# Subfunctionalized expression drives evolutionary retention of ribosomal protein paralogs in vertebrates

**DOI:** 10.1101/2022.05.03.490441

**Authors:** Adele Xu, Rut Molinuevo, Elisa Fazzari, Harrison Tom, Zijian Zhang, Julien Menendez, Kerriann M. Casey, Davide Ruggero, Lindsay Hinck, Jonathan K. Pritchard, Maria Barna

## Abstract

The formation of paralogs through gene duplication is a core evolutionary process. For paralogs that encode components of protein complexes such as the ribosome, a central question is whether they encode functionally distinct proteins, or whether they exist to maintain appropriate total expression of equivalent proteins. Here, we systematically tested evolutionary models of paralog function using the mammalian ribosomal protein paralogs *eS27 (Rps27)* and *eS27L (Rps27l)* as a case study. We first showed that *eS27* and *eS27L* have inversely correlated mRNA abundance across cell types, with the highest *eS27* in lymphocytes and the highest *eS27L* in mammary alveolar cells and hepatocytes. By endogenously tagging the eS27 and eS27L proteins, we demonstrated that eS27- and eS27L- ribosomes associate preferentially with different transcripts. Furthermore, we generated murine *eS27* and *eS27L* loss-of-function alleles that are homozygous lethal at different developmental stages. However, strikingly, we found that expressing eS27 protein from the endogenous eS27L locus, or vice versa, completely rescues loss-of-function lethality and yields mice with no detectable deficits. Together, these findings suggest that *eS27* and *eS27L* are evolutionarily retained because their subfunctionalized expression patterns render both genes necessary to achieve the requisite total expression of two equivalent proteins across cell types. Our work represents the most in-depth characterization of a mammalian ribosomal protein paralog to date and highlights the importance of considering both protein function and expression when investigating paralogs.

## Introduction

Gene duplication is a fundamental evolutionary process that expands the complexity and functional repertoire of genomes (Lynch & Conery, 2000; Nadeau & Sankoff, 1997; Ohno, 1970). A variety of evolutionary models have been proposed to explain why some gene duplicates, or paralogs, become fixed in a population and conserved over long evolutionary time scales (Conant & Wolfe, 2008; Innan & Kondrashov, 2010; Ohno, 1970; Prince & Pickett, 2002). For some genes, an increase in copy number may be directly advantageous (beneficial dosage increase) (Innan & Kondrashov, 2010; Kondrashov et al., 2002; Stark & Wahl, 1984). In other cases, duplication creates redundancy that relaxes the purifying selection on both paralogs. Such conditions may allow one paralog to evolve regulatory features for expressing in new contexts (neofunctionalized expression) (Force et al., 1999; Sidow, 1996), or to encode a new gene product (i.e. a neofunctionalized protein) (Ohno, 1970). Rather than allow new functions, redundancy can alternatively allow paralogs to partially degenerate. If *cis*- regulatory features are included in a duplication, a newly formed paralog pair should initially exhibit symmetric expression. Under the relaxed selection, each may accumulate regulatory mutations of similar effect and continue expressing symmetrically at a lower total level, but often one paralog eventually degenerates more than the other (asymmetric expression) or undergoes nonfunctionalization altogether (Lan & Pritchard, 2016; Lynch & Conery, 2000). It is also possible for two paralogs’ regulatory features to undergo complementary degeneration such that each becomes the major source of expression in a different subset of the ancestral contexts (subfunctionalized expression) (Force et al., 1999). Symmetric, moderately asymmetric, or subfunctionalized expression can promote retention of paralogs if they engage in dosage sharing, a situation in which both paralogs are needed to achieve the necessary total expression of the gene product (Force et al., 1999; Lan & Pritchard, 2016). Analogous to subfunctionalized expression, complementary degeneration of gene products (i.e. subfunctionalized proteins) can render both paralogs necessary to carry out the functions once performed by one ancestral gene product (Hughes, 1994). A final potential benefit of paralog retention is that one paralog may be able to compensate if the other paralog is disabled, thus conferring resilience against loss-of-function mutations (paralog buffering) (De Kegel & Ryan, 2019; Gu et al., 2003; Thompson et al., 2021).

Comparing theoretical models to experimental data on real-world paralogs is critical to understanding how genomes evolve and to predicting the effects of polymorphisms or therapeutic interventions involving paralogous genes. In terms of observable features in present-day paralogs, the above models fall into two categories: either the paralogs evolve to encode distinct gene products, or the expression of both paralogs becomes advantageous or necessary even if they encode similar gene products. Importantly, classic knockout, knockdown, and overexpression experiments cannot conclusively distinguish whether observed effects are caused by perturbing a function specific to one paralog’s gene product, or by altering total expression of two interchangeable gene products, or both. Instead, experimental designs that manipulate gene product characteristics without altering expression levels, or that isolate both gene products and their interactors from a source that expresses both paralogs, can help decouple the significance of expression level from that of divergent gene product characteristics. Editing an endogenous paralog gene to encode the gene product of the other paralog is an elegant approach of this nature that has occasionally been employed, as exemplified by previous work to substitute the exons of mouse *Hoxa3* for those of *Hoxd3* and vice versa (Greer et al., 2000), or to edit codons corresponding to differing residues between mouse actin paralogs (Patrinostro et al., 2018; Vedula et al., 2017). Rigorous experimental design was especially important in these cases because organisms are sensitive to the expression levels of Hox and actin genes (Greer et al., 2000; Vedula et al., 2017).

Sensitivity to expression level is also often encountered in genes encoding components of protein complexes, some of which require expression of all components at balanced dosages to achieve effective assembly (Papp et al., 2003; Taggart et al., 2020). However, it is also noteworthy that the functions of protein complexes can be modulated by incorporating alternative isoforms of their components (Antebi et al., 2017; Raices & D’Angelo, 2012). It is thus especially important to determine whether paralogs of protein complex components serve as necessary sources of expression, as functionally distinct proteins, or both. In particular, growing interest surrounds the paralogs of genes that encode the protein components of the ribosome, the ubiquitous macromolecular complex that catalyzes translation of all protein-coding transcripts (Genuth & Barna, 2018; Gerst, 2018; Komili et al., 2007; Topisirovic & Sonenberg, 2011). In mammals, each ribosome consists of 80 ribosomal proteins (RPs) and 4 ribosomal RNAs (rRNAs), which together form a 40S small subunit and a 60S large subunit that join during translation (Anger et al., 2013; Uechi et al., 2001). Each RP is encoded by a different genomic locus. Ten of these RP genes are known to exist as paralog pairs in the human genome, several of which are conserved among other mammals (Balasubramanian et al., 2009; Gupta & Warner, 2014; Nakao et al., 2004; Sugihara et al., 2010).

Two main categories of hypotheses could explain why some duplicated RP genes have been evolutionarily conserved. The first category posits that RP paralogs have diverged in their protein functions: they may have acquired distinct extraribosomal functions, which have been observed previously for certain RPs (Warner & McIntosh, 2009; Yong Zhang et al., 2017), or they may encode alternative RP isoforms that assemble to form ribosomes with different translational characteristics (Gerst, 2018). The latter possibility is especially intriguing, given that other variations in ribosome composition have been observed to affect protein synthesis or modulate the translation of specific genes (Genuth & Barna, 2018; Xue & Barna, 2012). Studies in yeast, in which 59 of the 79 RP genes exist as paralog pairs, have motivated speculation that functionally distinct RP paralogs are a major component of translational gene regulation through the formation of ribosomes that are heterogeneous in composition, a model known as the ‘ribosome code’ (Ghulam et al., 2020; Komili et al., 2007; Parenteau et al., 2015). Conversely, one could propose a second category of hypotheses that RP paralogs do not encode functionally distinct proteins, but are rather retained because their expression has become necessary. This possibility would be consistent with observations suggesting that, for paralogs in general, retention due to expression may be more common than divergence of protein function (Prince & Pickett, 2002). Such observations include systems-level analysis showing that paralogs found throughout fungal genomes rarely change gene ontologies or protein interaction networks (Wapinski et al., 2007), and experimental demonstrations for individual genes such as Hox paralogs that suggest equivalent protein function despite distinct expression patterns (Bruce et al., 2001; Greer et al., 2000).

In this work, we systematically examine these two categories of hypotheses in the case of the RP paralogs *eS27* and *eS27L* (known in classic RP nomenclature as *Rps27* and *Rps27l*, respectively). Previous work has shown that both *eS27* and *eS27L* are expressed in most tissues, and are incorporated into actively translating ribosomes (O’Donohue et al., 2010; Xiong et al., 2014). While no *eS27* knockout mouse has previously been described, homozygous *eS27L* knockout is lethal at early postnatal stages with increased apoptosis in hematopoietic organs (Xiong et al., 2014). *eS27* and *eS27L* are differentially regulated upon activation of the tumor suppressor *p53*, and have opposite feedback effects on the *p53*-*Mdm2* axis (He & Sun, 2007; Xiong et al., 2011, 2014). However, the study of *eS27* and *eS27L* is complicated by the consideration that impaired ribosome biogenesis due to perturbed dosage balance of other RPs has also been shown to activate *p53* via nucleolar stress or translation inhibition (Nicolas et al., 2016; Russo & Russo, 2017; Yanping Zhang & Lu, 2009). Thus, the knockout, knockdown, and overexpression experiments that have primarily been used to date could either reflect *eS27* and *eS27L’s* direct role in *p53* signaling, or the non-specific effects of perturbing ribosome biogenesis.

The challenges of interpreting the existing literature on *eS27* and *eS27L* informed our approach to investigating these paralogs without perturbing their expression. We first examined single-cell transcriptomics of normal mouse tissues and found previously unreported cell type-specific patterns of *eS27* and *eS27L* expression. We confirmed that the eS27 and eS27L proteins are incorporated into ribosomes, and used endogenously epitope-tagged eS27 and eS27L combined with immunoprecipitation and ribosome profiling to examine whether eS27L- and eS27L-ribosomes associate with different transcripts. We then examined the functions of *eS27* and *eS27L* at an organismal level, by first generating loss-of-function alleles for each gene and, most importantly, by generating ‘homogenized’ mouse lines in which the *eS27* locus has been edited to encode the eS27L protein, or vice versa. Finally, we performed a detailed characterization of the homogenized mouse lines with particular attention to the cell types that preferentially express one paralog, and also examined the effects of paralog homogenization on aging and response to genotoxic stress.

In sum, investigating the functions of RP paralogs is valuable for elucidating the evolutionary fates of paralogs that encode components of protein complexes, and may shed light on the potential role of paralogs in the ribosome code. Such work demands careful distinctions between the possibility that paralogs encode functionally distinct proteins, as opposed to the possibility that they provide expression of similar proteins. With close attention to these challenges, we present here the most in- depth profiling of a mammalian RP paralog to date, in which we have leveraged single-cell transcriptomics, molecular assays, and mouse genetics to comprehensively examine the above hypotheses regarding paralog function in the case of *eS27* and *eS27L*.

## Results

### *eS27* and *eS27L* are vertebrate ohnologs encoding highly conserved proteins

To guide our functional analysis of *eS27* and *eS27L*, we first considered their probable evolutionary trajectory by examining their chromosomal arrangement and sequence homology. In general, several lines of evidence can support whether a duplicated gene arose through DNA-based events such as DNA transposition, tandem duplication, segmental duplication, or whole-genome duplication (WGD); or through retrotransposition of RNA (reviewed in (Graur & Li, 1997)). In human and mouse, *eS27* and *eS27L* are located on separate chromosomes. Unlike the approximately 2,000 processed RP pseudogenes resulting from retrotransposition in these genomes (Balasubramanian et al., 2009), the canonical transcripts for *eS27* and *eS27L* each contain three introns. While their nucleotide sequences are highly divergent, almost all substitutions are synonymous. Thus, the protein sequences only differ by three out of 84 residues, which are near the N-terminus: K5R, P12L, and R17K. Of the vertebrate genomes examined, all had at least two loci encoding proteins resembling *eS27* and *eS27L*, and most had the same differing residues as human and mouse (Figure 1A, Supplementary Table 1). Meanwhile, the non-vertebrate genomes examined each only had one locus encoding a protein resembling eS27. Thus, *eS27* gene duplication most likely occurred through a DNA- based event in an ancestor common to most vertebrates. Chronologically, this coincides with the ‘1R’ and ‘2R’ WGDs that occurred early in vertebrate evolution at least 400 million years ago (Dehal & Boore, 2005; Ohno, 1970). Indeed, in systematic efforts to map 1R/2R remnants based on phylogeny and large-scale synteny (Makino & McLysaght, 2010; Sacerdot et al., 2018; Singh & Isambert, 2020), *eS27* and *eS27L* are consistently identified among the ‘ohnolog’ gene duplicates that still comprise 25- 35% of present-day vertebrate genes.

**Figure 1:**
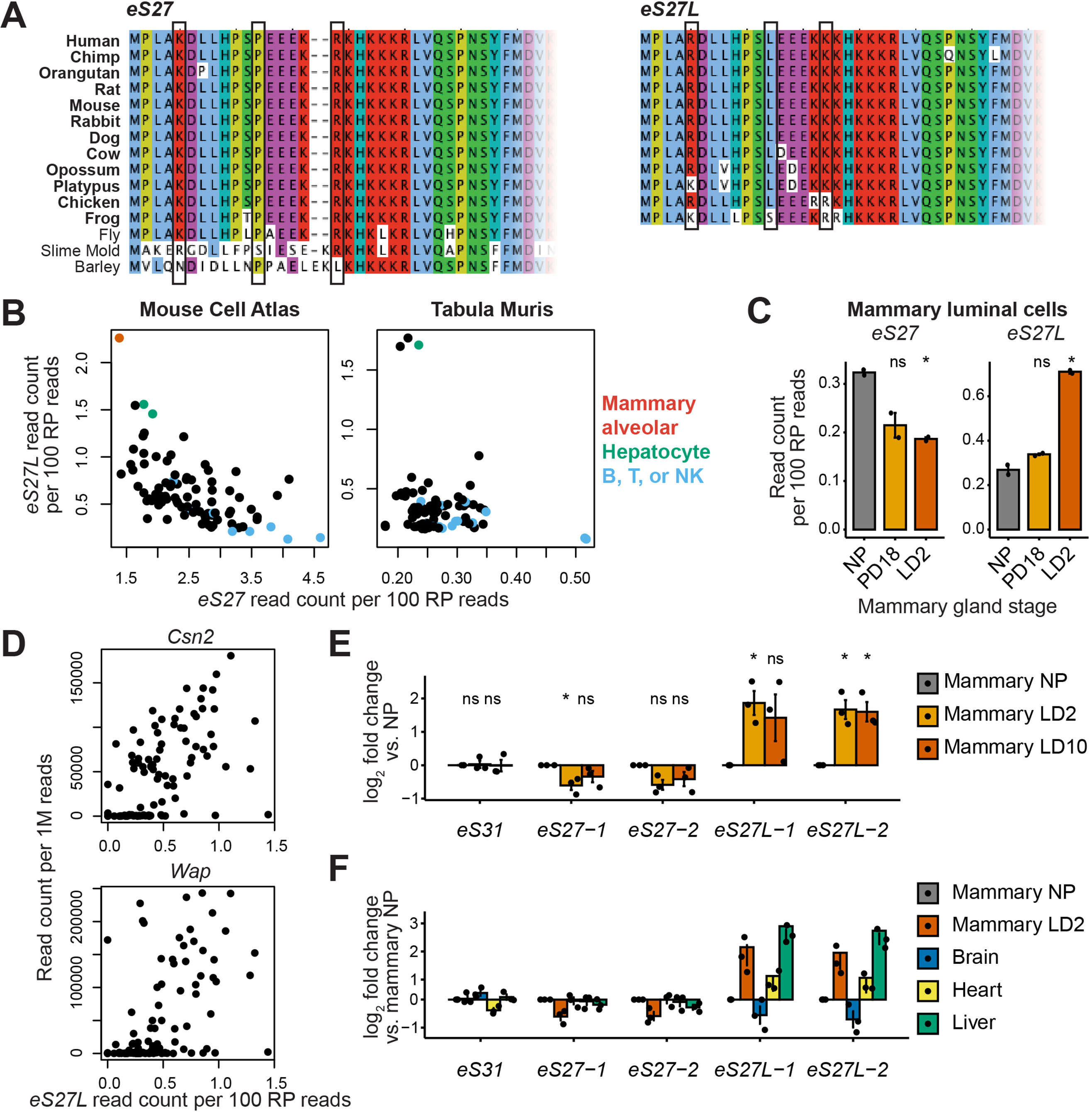
*eS27* and *eS27L* protein sequence conservation and cell type-specific mRNA abundance. **(A)** Multi-species alignment of eS27 and eS27L N-terminal protein sequences among vertebrates (bolded) and invertebrates. Full species names and Uniprot IDs are listed in Supplementary Table 1. The three residues that differ between eS27 and eS27L in human and mouse are boxed. **(B)** *eS27* and *eS27L* scRNA-seq values across cell types from the Mouse Cell Atlas (Han et al., 2018; Pearson’s r = -0.58, p = 4.4e-09) and Tabula Muris (Tabula Muris Consortium et al., 2018; Pearson’s r = -0.23, p = 0.024). See also Supplementary Figure 1. **(C)** *eS27* and *eS27L* RNA-seq values (Fu et al., 2015) in flow-sorted mammary luminal cells from nulliparous (NP), pregnancy day 18.5 (PD18), and lactation day 2 (LD2) mice. n = 2 biological replicates (individual animals). Significance compared to NP was assessed by t-test. *eS27* LD2: p=0.0049; *eS27L* LD2: p=0.016. “ns” indicates p > 0.05. **(D)** Correlation between *eS27L* and milk protein transcripts in scRNA-seq (Bach et al., 2017) from alveolar cells in lactating mammary glands (‘Avd-L’ cells as termed by Bach et al.). *Csn2*: Spearman’s ρ = 0.63, p = 5.7e-13. *Wap*: Spearman’s ρ = 0.56, p = 6.7e-10. **(E)** RT-qPCR of *eS27*, *eS27L*, and other ribosomal protein (RP) genes in mam- mary glands of nulliparous and lactating female mice. Values are normalized to RP *eS6* and are shown as log fold differences over NP. For *eS27* and *eS27L*, two independent primer sets (“-1” and “-2”) were used. n = 3 biological replicates. Significance compared to NP was assessed by t-test. eS27-1 LD2: p = 0.050; eS27L-1 LD2: p = 0.035; eS27L-2 LD2: p = 0.028; eS27L-2 LD10: p = 0.032. “ns” indicates p > 0.05. **(F)** RT-qPCR of *eS27*, *eS27L*, and other RP genes in mammary glands, heart, liver, and brain of LD2 female mice. Values are normalized to RP *eS6* and are shown as log fold differences over NP mammary gland. n = 3 biological replicates.

The probable evolutionary trajectory of eS27 and eS27L should also be considered in relation to other RPs: in fungi, genes encoding protein complexes such as the ribosome often remain duplicated after WGD but rarely duplicate on an individual basis. This observation supports a model that assumes dosage balance is required between components of a protein complex, such that duplicating one gene would cause a deleterious imbalance but simultaneously duplicating all the components would maintain balance and favor initial retention of the resulting paralogs (Papp et al., 2003; Wapinski et al., 2007).

Notably, most RPs in present-day mammalian genomes are encoded by a single gene; there are only ten known human RP paralog pairs, several of which have hallmarks of tandem duplication or retrotransposition rather than WGD (Gupta & Warner, 2014). Two evolutionary trajectories are thus possible: either all RP genes were duplicated via WGD and *eS27(L)* is one of the few that did not eventually revert to a single gene through nonfunctionalization of one duplicate, or a smaller-scale duplication occurred for eS27 alone and survived despite initial excess dosage. Based on the above evidence that *eS27* and *eS27L* are vertebrate-specific and are located within large mutually syntenic regions, we consider 1R/2R WGD to be the more likely origin.

Based on these evolutionary features of eS27 and eS27L, we can preliminarily assess the relevance of the proposed paralog retention models. Given the evidence from fungi that RP genes are subject to dosage balance constraints, beneficial dosage increase and neofunctionalized expression are less likely models of paralog retention in this case. However, the other proposed models all remain plausible. In contrast to tandem duplications in which both paralogs reside near the same regulatory features (Lan & Pritchard, 2016), or transpositions that introduce a duplicate into a completely different regulatory context, WGD produces duplicates on separate chromosomes with initially identical regulatory features that may then diverge. The protein sequences of eS27 and eS27L have remained highly conserved over their long evolutionary history, suggesting that they still perform related molecular roles in the cell and could participate in dosage sharing or paralog buffering. On the other hand, the three differing residues could confer partially distinct protein functions by affecting protein structure or post-translational modifications. Thus, we proceeded to compare both the expression patterns and the protein characteristics of *eS27* and *eS27L*.

### *eS27* and *eS27L* mRNA expression are cell type-dependent

We next leveraged publicly available single-cell RNA-seq datasets to compare expression levels of *eS27*, *eS27L*, and other RP genes in previously unexamined primary cell types. Different cell types express RP genes at different levels, likely reflecting the ribosome production rate needed to accommodate each cell type’s translational or proliferative demands. However, transcript abundance among the core RP genes is highly correlated across cell types (Supplementary Figure 1A), which is consistent with the theory that components of a protein complex must have balanced expression for effective assembly (Guimaraes & Zavolan, 2016; Papp et al., 2003). Thus, for each cell, we used the summed transcript abundances of all RP genes to normalize each RP gene’s expression level by the cell type’s ribosome production rate, allowing us to identify RP genes with disproportionately high or low expression relative to other RP genes in a cell type.

By analyzing single-cell data from annotated cell types across dozens of mouse tissues in the Mouse Cell Atlas (Han et al., 2018), we first were able to corroborate the previously reported tissue- specific expression patterns of several other RP paralogs (Supplementary Figure 1B) (Chaillou et al., 2016; Guimaraes & Zavolan, 2016; Jiang et al., 2017; Sugihara et al., 2010; Wong et al., 2014).

Interestingly, we also found that *eS27* and *eS27L* mRNA levels are inversely correlated across cell types (Figure 1B). The highest *eS27L:eS27* ratios were found in mammary alveolar cells and hepatocytes; the lowest were found in a subset of B, T, and NK lymphocytes. A similar expression pattern was detected among FACS-isolated hepatocytes and lymphocytes collected in Tabula Muris (Tabula Muris Consortium et al., 2018), a separately constructed large dataset of mouse single-cell RNA-seq samples (Figure 1B).

Mammary alveolar cells are a transient cell type that arise rapidly from luminal progenitor cells in female mammary glands during pregnancy and lactation to secrete milk components (Macias & Hinck, 2012). Because lactating mammary glands were not collected in Tabula Muris, we corroborated the Mouse Cell Atlas findings by analyzing a single-cell RNA-seq dataset of mammary development timepoints (Bach et al., 2017), and a bulk RNA-seq dataset of sorted mouse mammary cell types (Fu et al., 2015). We found that eS27L mRNA levels increase in luminal cells between pregnancy and lactation, while eS27 levels diminish (Figure 1C). Among mature alveolar cells, single cells expressing the highest levels of milk protein transcripts such as *Csn2* and *Wap* also had the highest *eS27L* expression (Figure 1D). Indeed, these transcripts correlated more strongly with *eS27L* expression than almost all others in the transcriptome (Supplementary Table 2). Notably, these datasets were collected from mice of different strains, suggesting a generalizable trend.

To confirm these *in silico* findings, we performed RT-qPCR for *eS27* and *eS27L* on bulk mouse mammary tissue from nulliparous and lactating females (Figure 1E). Liver, heart, and brain samples from the same animals were also analyzed (Figure 1F). Liver and lactating mammary gland had higher levels of *eS27L* and lower levels of *eS27* than nulliparous mammary gland, heart, or brain. Ribosomal protein *eS6* (*Rps6*) was used to normalize for the ribosome production rate of each tissue. Ribosomal protein *eS31* (*Rps27a*) was included as an example of an RP whose transcript abundance does not change during lactation when normalized by *eS6*.

It has previously been reported that *eS27L* is transcriptionally upregulated by p53 via two p53- binding elements within the *eS27L* genomic locus, and that p53 also downregulates *eS27* expression (He & Sun, 2007; Li et al., 2007; Xiong et al., 2011, 2014). To determine whether p53 activity and *eS27* or *eS27L* expression may be correlated in the mammary gland, we performed RT-qPCR for *p53*, *eS27*, *eS27L*, and other RP transcripts on bulk mammary tissue at nulliparous, pregnant, and lactating timepoints (Figure 2A). *p53* transcript abundance is stable during early-to-mid pregnancy, whereas *eS27* and *eS27L* both decrease within the first 6 days of pregnancy. *p53* transcript abundance then decreases in the latter half of pregnancy, whereas *eS27L* increases and *eS27* remains decreased around lactation day 2 (LD2). Recognizing that p53 activity can be modulated through post-transcriptional mechanisms not reflected by *p53* mRNA abundance (Ashcroft & Vousden, 1999; Meek & Anderson, 2009), we again used a single-cell RNA-seq dataset of mammary gland timepoints (Bach et al., 2017) to compare *eS27L* expression against the expression of other genes that are transcriptionally activated by p53 (Figure 2B). Among differentiated mammary alveolar cells, we found that *eS27L* expression is inversely correlated with mRNA abundance of other p53-upregulated genes, suggesting that single cells with high *eS27L* expression do not have increased p53 activity. These findings therefore suggest that differences in *eS27* and *eS27L* expression across cell types are not primarily driven by p53 activity.

**Figure 2:**
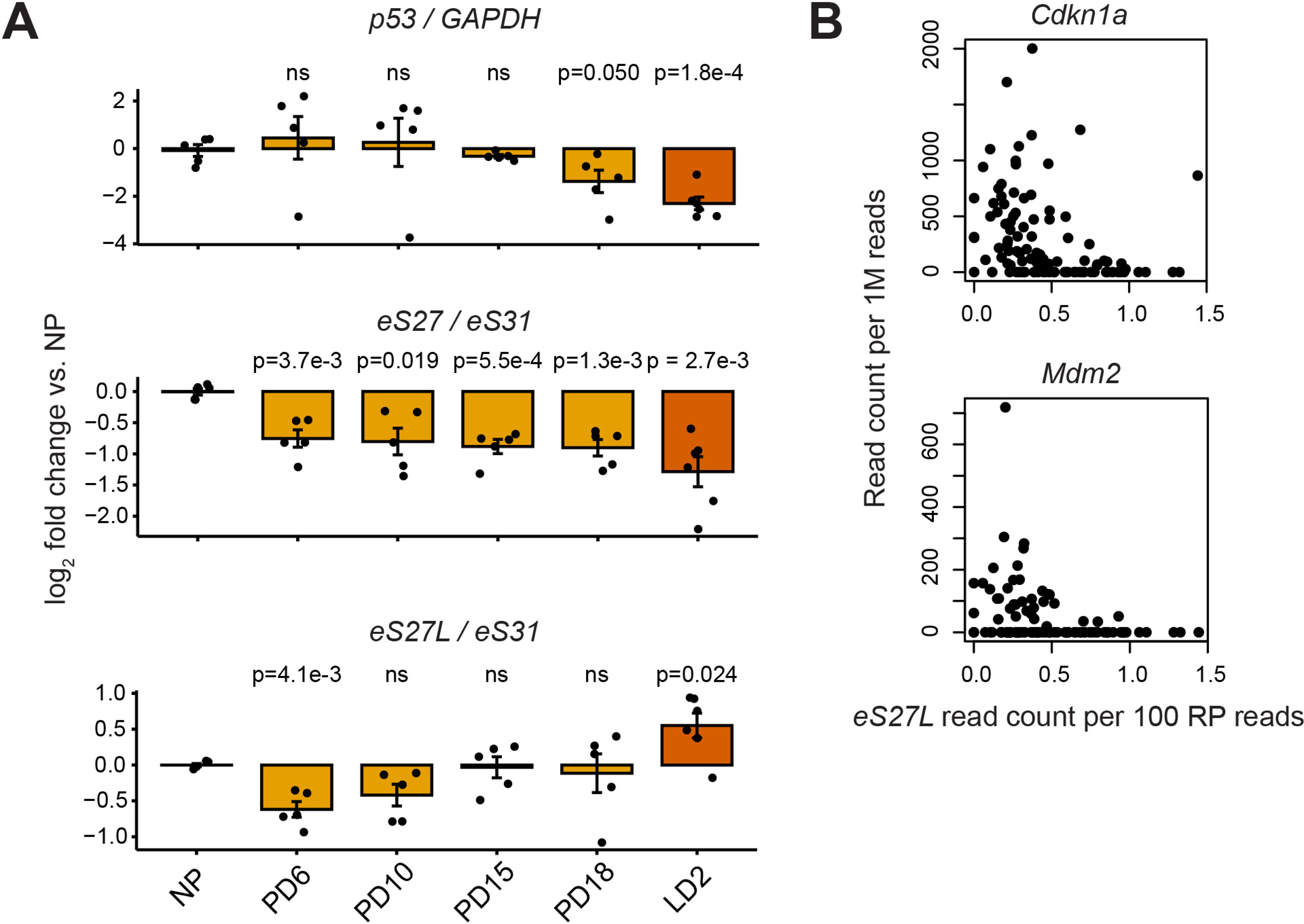
Comparison of *eS27* and *eS27L* mRNA abundance versus *p53* expression and activity. **(A)** RT-qPCR of *eS27*, *eS27L*, and *p53* in mammary glands at nulliparous (NP), pregnancy day 6-18 (PD6-18), and lactation day 2 (LD2) timepoints. *eS27* and *eS27L* are normalized by ribosomal protein (RP) *eS31*. *p53* is normalized by *GAPDH*. n = 5-6 biological replicates (individual animals) per timepoint. Significance versus NP was assessed by t-test. “ns” indicates p > 0.05. **(B)** Correlation between *eS27L* and other targets of p53 transcriptional activation in scRNA-seq of alveolar cells from lactating mammary glands (Bach et al., 2017). *Cdkn1a*: Spearman’s ρ = -0.51, p = 3.7e-8. *Mdm2*: Spearman’s ρ= -0.37, p = 1.2e-4.

Our findings that *eS27* and *eS27L* have inversely correlated mRNA abundance across cell types suggest that *eS27* and *eS27L* have complementary expression patterns, a hallmark of subfunctionalized expression. While *eS27* and *eS27L* have previously been included in bulk transcriptomic analysis of RPs (Guimaraes & Zavolan, 2016; Gupta & Warner, 2014; He & Sun, 2007), our observations were facilitated by including lactating mammary gland samples and achieving single-cell resolution of cell types. Importantly, dosage sharing via subfunctionalized expression could alone explain the retention of these paralogs through a duplication-degeneration-complementation (DDC) process (Force et al., 1999): whereas the ancestral *eS27* gene may have been expressed widely, the two paralogs may have lost complementary sets of regulatory elements until both were needed to maintain the appropriate expression level across all cell types. If *eS27* and *eS27L* are indeed remnants of a WGD as discussed in the previous section, DDC of *eS27* and *eS27L* may have happened in concert with the other RP paralog pairs’ reversion to single genes.

### eS27 and eS27L-ribosomes differentially associate with cell cycle-related mRNAs

Having established their distinct expression patterns, it remained to be explored whether the two paralogs could not only have different expression but also encode functionally different proteins, which has been observed with other paralogs (Conant & Wolfe, 2006). Using mouse embryonic stem cells (mESCs) as a primary cell line amenable to genetic editing, we first confirmed whether eS27 and eS27L are incorporated into actively translating ribosomes. mESC lysate was fractionated over a sucrose density gradient to separate mRNAs based on the number of ribosomes bound to them. Using antibodies specific to each paralog (Supplementary Figure 2A), we found that eS27 and eS27L are both detectable among fractions corresponding to actively translating ribosomes (polysomes), fractions corresponding to single ribosomes (80S), and fractions representing the 40S small subunit. eS27 and eS27L are not detectable in early fractions where most extra-ribosomal proteins are found (Figure 3A). In the structure of the human ribosome (Natchiar et al., 2017), eS27 is incorporated into the 40S subunit such that the three paralog-specific amino acid positions are solvent-accessible (Figure 3B) and could modulate interactions with other molecules.

**Figure 3:**
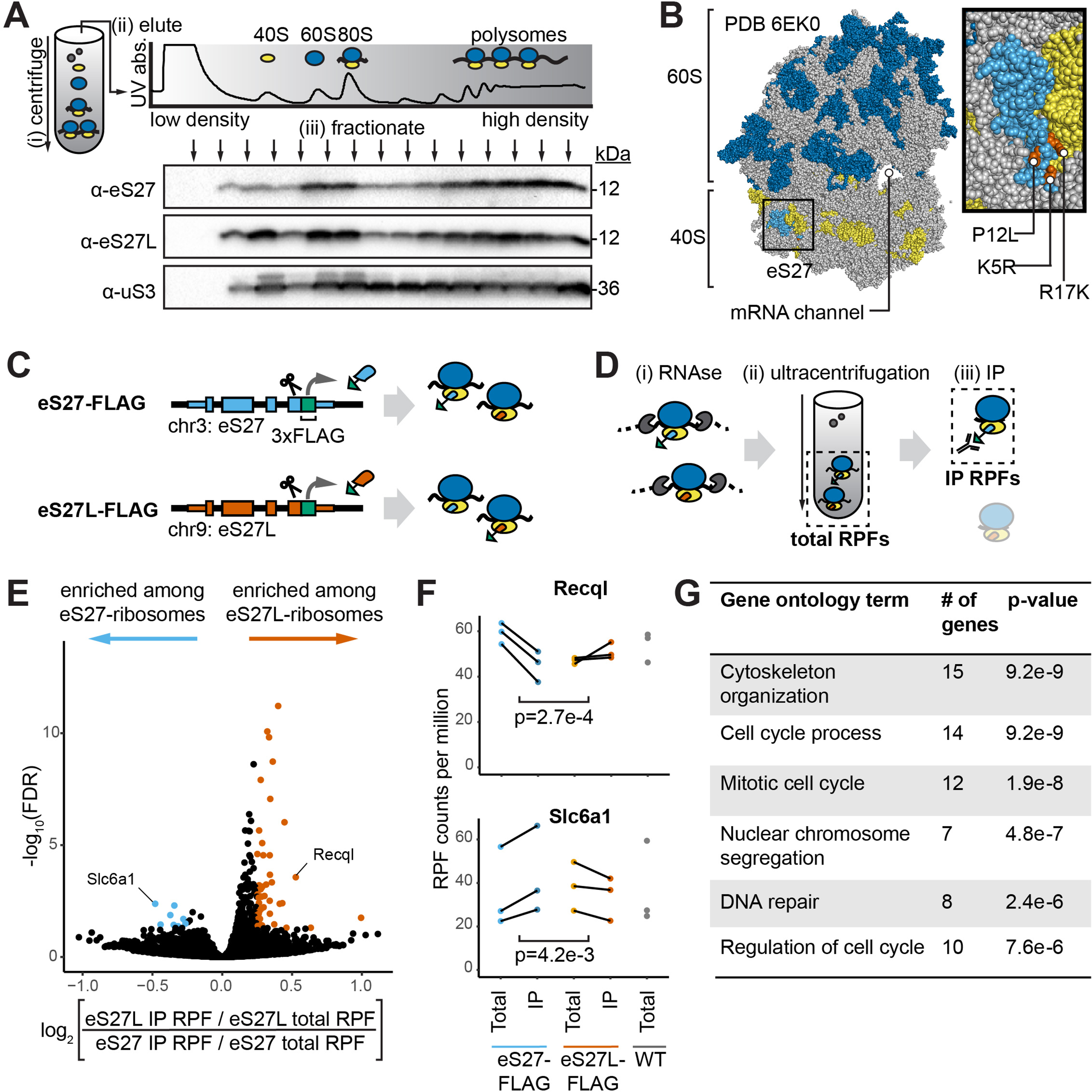
Paralog-specific ribosome profiling reveals preferential association of eS27L- ribosomes with cell cycle-related mRNAs. **(A)** Fractionation of mouse embryonic stem cell (mESC) lysate by density to separate non-ribosomal proteins, ribosome 40S and 60S subunits, mRNAs bound by one ribosome (80S), and mRNAs bound by multiple ribosomes (polysomes). See Figure 3--Source Data 1-3. **(B)** Ribosome structure showing the 60S subunit (with 28S rRNA in dark blue), 40S subunit (with 18S rRNA in yellow), and eS27 protein (light blue) within the 40S. Orange denotes residues that differ between eS27 and eS27L. Gray denotes other RPs. **(C)** CRISPR-mediated insertion of 3xFLAG epitope tag C-terminally at the endogenous eS27 and eS27L genomic loci in mESCs. **(D)** FLAG-IP ribosome profiling of paralog-containing ribosomes. i) Cell lysate is treated with RNAse to digest mRNA sequences that are not protected by bound ribosomes. ii) Ribosomes and associated ribosome-protected fragments (RPFs) are collected by ultracentrifugation. These are termed total RPFs. iii) anti- FLAG immunoprecipitation (IP) is performed on total ribosomes. IP RPFs are eluted. **(E)** Comparison of RPFs enriched after IP in eS27L-FLAG versus eS27-FLAG mESCs. **(F)** Examples of genes differentially enriched upon eS27-FLAG or eS27L-FLAG pulldown. Significance was assessed using an empirical Bayes method for detecting differential expres- sion, applied to a multilevel linear model to find genes whose RPF abundance is differentially affected by IP from eS27- and eS27L-FLAG mESCs. n =3 biological replicates (independent mESC clones). Multiple-hypothesis-corrected false discovery rates (FDR) are shown. **(G)** Top enriched gene ontology terms for RPFs that preferentially associate with eS27L-ribosomes. See also Supplementary Figures 2 and 3.

To compare the molecular interactions of eS27- and eS27L-ribosomes, we devised a strategy to isolate the two ribosome populations from *in vitro* cultured mouse cells and focused on comparing the mRNAs with which they associate. To enable isolation of eS27- and eS27L-ribosomes without overexpressing an exogenous epitope-tagged construct, we used CRISPR to homozygously insert 3xFLAG epitope tags immediately preceding the stop codon at the *eS27* and *eS27L* loci (Figure 3C).

This approach yielded three independently selected clones (biological replicates) of eS27-FLAG mESCs and three of eS27L-FLAG mESCs. eS27- and eS27L-FLAG proteins are expressed at comparable levels to the untagged proteins and are incorporated into actively translating ribosomes (Supplementary Figure 2B-C). We confirmed by Western blot that anti-FLAG immunoprecipitation (FLAG-IP) efficiently enriches for eS27- and eS27L-ribosomes from the eS27-FLAG and eS27L-FLAG mESCs, respectively (Supplementary Figure 2D). Minimal eS27 is detected when targeting eS27L for pulldown and vice versa, confirming that a ribosome does not simultaneously contain eS27 and eS27L and that we did not isolate undigested polysomes containing multiple ribosomes (Supplementary Figure 2D). We then performed ribosome profiling on the total ribosome population in each line and on the paralog-containing ribosomes isolated via FLAG-IP (Figure 3D). Ribosome profiling identifies mRNA regions occupied by ribosomes, using RNAse digestion to enable sequencing of ribosome-protected fragments of mRNA (RPFs) (Ingolia et al., 2009). The total RPFs from the two FLAG-tagged cell lines did not contain mRNAs that differed significantly in abundance, either in comparison between the eS27- FLAG line and eS27L-FLAG line, or in comparison to a passage-matched wild-type (WT) control line (Supplementary Figure 2E). This confirms that FLAG-tagging did not alter the landscape of normally translated mRNAs. We then compared the IP RPFs from both cell lines to each other, using the total RPFs from each clone to normalize for its overall translational landscape (Figure 3E-F). After excluding a *Tmod3* transcript as a likely artifact of FLAG-IP (Supplementary Text, Supplementary Figure 3), we identified 8 transcripts enriched among eS27-ribosomes, and 46 transcripts enriched among eS27L- ribosomes (absolute value of log2 fold change > 0.25, false discovery rate < 0.05) (Supplementary Table 3). Among the latter set of transcripts, gene ontology terms associated with cell cycle processes were enriched (Figure 3G). These findings intriguingly demonstrated that eS27- and eS27L-ribosomes associate differently with specific mRNAs.

Importantly, our experimental design minimized the possibility that these results reflect translational changes due to genetically editing the cell lines: no knockdown, knockouts, or overexpression were used, and the IP RPF transcript abundances were normalized by the total RPF transcript abundances for each clone. A critical consideration, however, is that this approach demonstrates a correlation between eS27L incorporation and ribosome association with cell cycle-related transcripts, but it does not directly demonstrate that eS27L causes ribosomes to preferentially bind these transcripts. Nevertheless, to our knowledge, this is the first instance of comparing RP paralogs by endogenously epitope-tagging each paralog and performing IP-ribosome profiling. To definitively compare the functions of the two proteins, we next turned to *in vivo* approaches.

### *eS27* and *eS27L* homozygous knockouts are lethal at different developmental stages

To compare *eS27* and *eS27L in vivo*, we first examined the organism-level effects of knocking out each paralog. We generated mice harboring two truncation alleles at the endogenous *eS27* and *eS27L* loci: *eS27^exon2del^*, in which the splicing junctions flanking exon 2 of *eS27* are deleted; and *eS27L^exon2del^*, which harbors a 320 bp deletion in *eS27L* that spans exon 2 and part of the subsequent intron (Figure 4A). Similar to a previously described *eS27L* gene-trapped loss-of-function mouse model (here termed *eS27L^GT^*) (Xiong et al., 2014), *eS27L^exon2del / +^* males and females are viable and fertile, but *eS27L^exon2del / exon2del^* mice are observed at lower-than-expected frequencies in crosses of *eS27L^exon2del / +^* males and females (Figure 4B). These *eS27L^exon2del / exon2del^* offspring died or needed to be euthanized due to animal distress by postnatal day 13-17 (P13-17). Interestingly, *eS27^exon2del / +^* mice were also viable, but no *eS27^exon2del / exon2del^* offspring were recovered from crosses of *eS27^exon2del / +^* males and females (Figure 4B). When we dissected embryonic day 8.5 (e8.5) embryos from *eS27^exon2del / +^* x *eS27^exon2del / +^* crosses, only one *eS27^exon2del / exon2del^* specimen out of 35 dissected embryos was recovered. This *eS27^exon2del / exon2del^* embryo was severely delayed in development compared to littermates (Figure 4C). Thus, impaired expression from the *eS27* locus impacts viability at significantly earlier embryonic stages than impaired expression from the *eS27L* locus.

**Figure 4:**
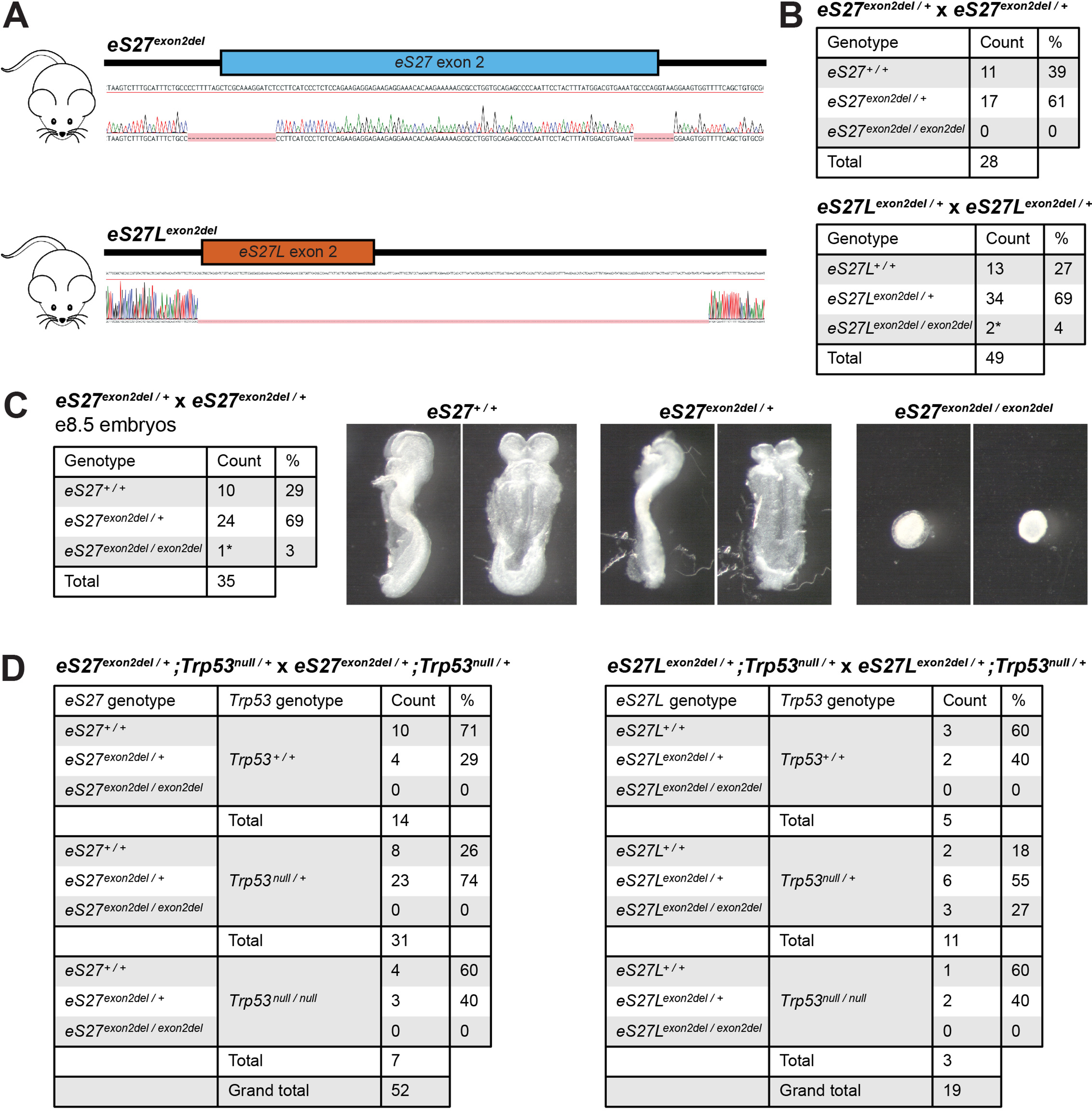
Truncation alleles of *eS27* and *eS27L* are homozygous lethal at embryonic and early postnatal stages respectively. **(A)** Sanger sequencing of *eS27^exon2del^* and *eS27L^exon2del^* alleles. **(B)** Genotype ratios among live offspring of *eS27^exon2del / +^* and *eS27L^exon2del / +^* heterozygous crosses at postnatal day 13 (P13). * = animals that died or needed to be euthanized at P13-17. **(C)** Genotype ratios and photographs of dissected embryos of *eS27^exon2del / +^* heterozygous crosses at embryonic day 8.5 (e8.5). * = embryo with severely delayed development. **(D)** Genotype ratios among live offspring of *eS27^exon2del / +^;Trp53^null / +^* double heterozygous crosses and *eS27L^exon2del / +^;Trp53^null / +^* double heterozygous crosses at P13.

The phenotypes of these *eS27^exon2del^* and *eS27L^exon2del^* loss-of-function alleles yield several preliminary insights on the possible roles of these RP paralogs *in vivo*. The fact that *eS27* and *eS27L* loss-of-function alleles are each lethal in homozygosity, even with both alleles of the other paralog intact, disfavors the hypothesis that *eS27* and *eS27L* engage in paralog buffering. There is, however, a difference in the timing of lethality that could be consistent with the two proteins having distinct functions; under such a model, it would appear that eS27 protein has a critical function in very early developmental processes, while the functions of eS27L protein are dispensable until postnatal stages. However, divergent protein function is not the only possible explanation: the timing difference could also be explained by early reliance on expression from the *eS27* locus, whereas this dependence later shifts to expression of an equivalent protein from the *eS27L* locus. If the regulatory characteristics of *eS27* and *eS27L* are sufficiently dissimilar, it may be impossible for one paralog to compensate for the other’s loss of function in specific cell types.

Previous work has shown that the early postnatal lethality of *eS27L^GT/GT^* can be rescued on a *p53* loss-of-function (*p53^LOF/+^* or *p53^LOF/LOF^*) background. This finding was attributed to a model in which *eS27L* depletion impairs ribosome biogenesis and thus causes accumulation of unincorporated RPs, which increase p53 activity by blocking its degradation by Mdm2. The increased p53 triggers apoptosis in hematopoietic tissues (Xiong et al., 2014). We thus crossed the *eS27^exon2del^* and *eS27L^exon2del^* mice to *Trp53^null^* mice. We indeed found that *eS27L^exon2del / exon2del^;Trp53^null / +^* mice are viable. However, no *eS27^exon2del / exon2del^;Trp53^null / +^* or *eS27^exon2del / exon2del^;Trp53^null / null^* offspring were recovered from crossing *eS27^exon2del / +^;Trp53^null / +^* males and females (Figure 4D). These results suggest that disabling expression from the *eS27* locus disrupts ribosome biogenesis either to a greater degree than *eS27L* that cannot be mitigated by *p53* depletion, or through a non-*p53*-mediated mechanism.

### eS27 and eS27L proteins are functionally interchangeable across all examined murine tissues

Having demonstrated that expression of both *eS27* and *eS27L* are essential to development at different stages, we next asked whether eS27 protein can rescue loss of eS27L protein, and vice versa, at the whole-organism level. This test is critical to understanding whether the two paralogs’ gene products are functionally interchangeable. In order to do so rigorously, it was ideal to express the swapped protein sequences from the endogenous genomic loci with minimal perturbation to the regulatory contexts. Using CRISPR in mouse embryos, we minimally edited the endogenous genomic loci for *eS27* and *eS27L* to encode the protein sequence of the other paralog. This yielded two novel mouse alleles: *eS27^eS27L^*, which expresses eS27L protein from the *eS27* locus; and *eS27L^eS27^*, which expresses eS27 protein from the *eS27L* locus (Figure 5A-B).

**Figure 5:**
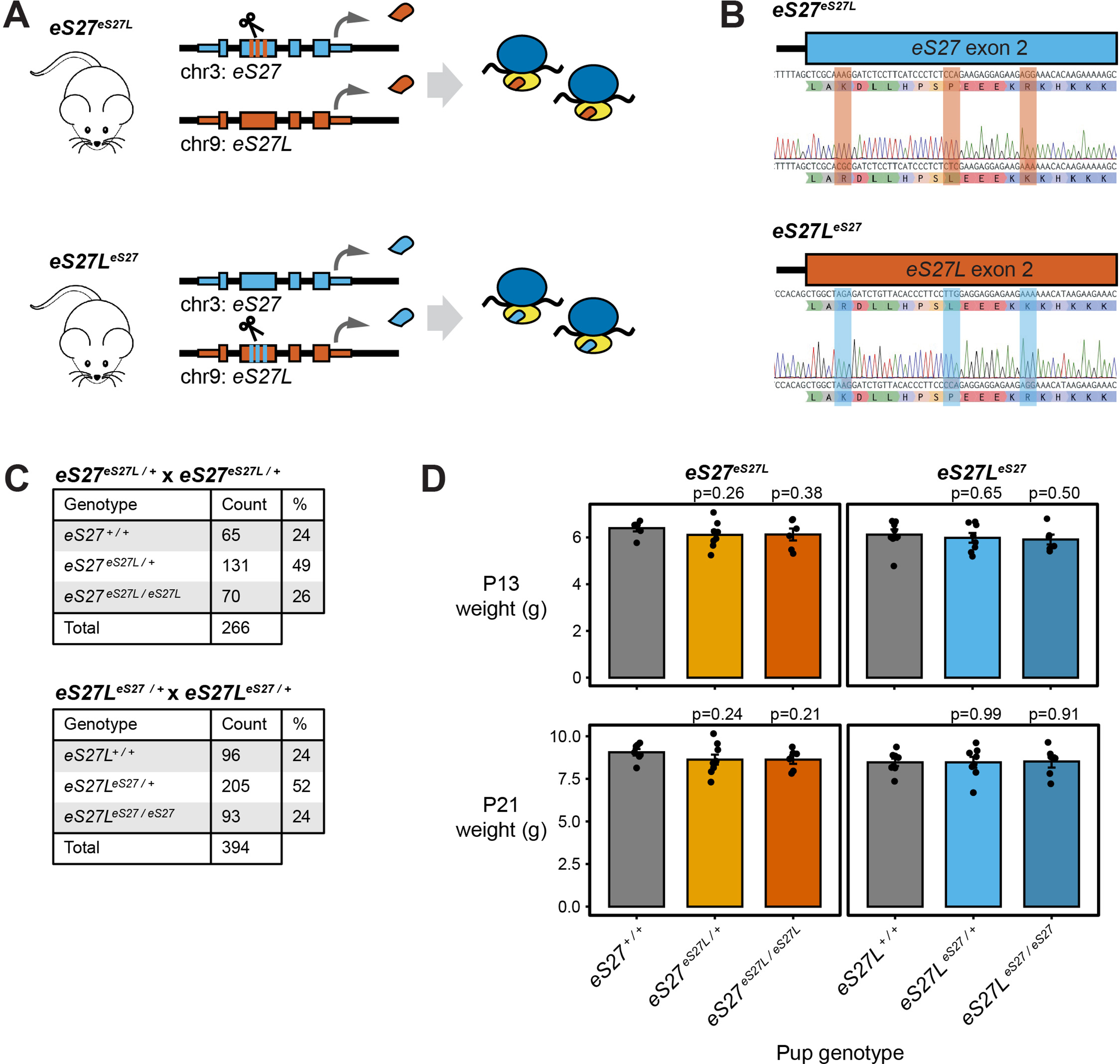
Homogenized *eS27* and *eS27L* alleles in mice exhibit normal genotype ratios and early development. **(A)** CRISPR editing to generate *eS27^eS27L^* and *eS27L^eS27^* homogenized mice. **(B)** Sanger sequencing of *eS27^eS27L^* and *eS27L^eS27^* homogenized mouse alleles **(C)** Genotype ratios among live offspring of *eS27^eS27L / +^* and *eS27L^eS27 / +^* heterozygous crosses at P13. **(D)** Pup weights at P13 and P21, grouped by pup genotype. Each data point represents the average weight among pups of the indicated genotype within a litter. Only pups from the first litters born to *eS27^eS27L / +^* and *eS27L^eS27 / +^* dams are included. n = 6-9 litters per genotype. Significance versus *eS27 ^+ / +^* and *eS27L^+ / +^*, respectively, was assessed by t-test. See also Supplementary Figure 4.

To assess the organism-wide impact of substituting the eS27 protein sequence with eS27L or vice versa, we performed a detailed characterization of *eS27^eS27L^* and *eS27L^eS27^* mice. Remarkably, heterozygous crosses for both alleles (*eS27^eS27L / +^* x *eS27^eS27L / +^*, and *eS27L^eS27 / +^* x *eS27L^eS27 / +^*) resulted in normal genotype frequencies among offspring of both sexes (Figure 5C). This demonstrates that eS27 completely rescues the early lethality observed upon homozygous truncation of eS27L, and vice versa. Later organism fitness was also rescued: pups of all genotypes gained weight at similar rates (Figure 5D), and male and female heterozygous and homozygous mice are viable to at least one year of age and are fertile. A detailed necropsy of homozygous 10-16-week-old males and age-matched wild-type controls, performed by a veterinary pathologist blinded to specimen genotype, revealed no clinically significant differences in gross organ weight, gross organ morphology, or tissue histology for either line (Supplementary Table 4) upon examination of neurological, cardiovascular, respiratory, gastrointestinal, genitourinary, lymphatic, endocrine, hematopoietic, integumentary, and musculoskeletal tissues.

Given the cell type-specific patterns of *eS27* and *eS27L* mRNA levels that we have described above, it was important to explore whether a functional difference between the eS27 and eS27L proteins might only be apparent in tissues that preferentially express one paralog or the other. The top cell types of interest included mammary alveolar cells and hepatocytes, which have a high *eS27L:eS27* ratio; and B, T, and NK cells, which have a low ratio (Figure 1B). We therefore devoted additional effort to characterizing the mammary gland, liver, and hematopoietic organs. A liver panel and complete blood count were performed on the 10-16-week-old male necropsy specimens to survey for anomalies in liver function or hematopoiesis. These yielded no clinically significant differences between genotypes (Figure 6A, Supplementary Table 5). To assess the mammary gland, we focused on the pregnancy and lactation stages because precursors to alveolar cells emerge and mature during these stages (Macias & Hinck, 2012), and because we had detected fluctuations in *eS27* and *eS27L* mRNA abundance in bulk tissues from these timepoints relative to nulliparous samples (Figure 1C-F). We first tracked pup weight gain as a metric of mammary gland function. Nulliparous littermate-matched females at 8 weeks of age were housed with stud males and separated from the males before parturition. Since genetically manipulated mouse lines can exhibit lactation failure at a range of timepoints (Palmer et al., 2006), litter size and pup weight were assessed at P4, P13, and P21 (Figure 6B). No statistically significant differences in pup weight gain were detected between any of the maternal genotypes. There were also minimal differences in litter size or maternal age at parturition (Supplementary Figure 4).

**Figure 6:**
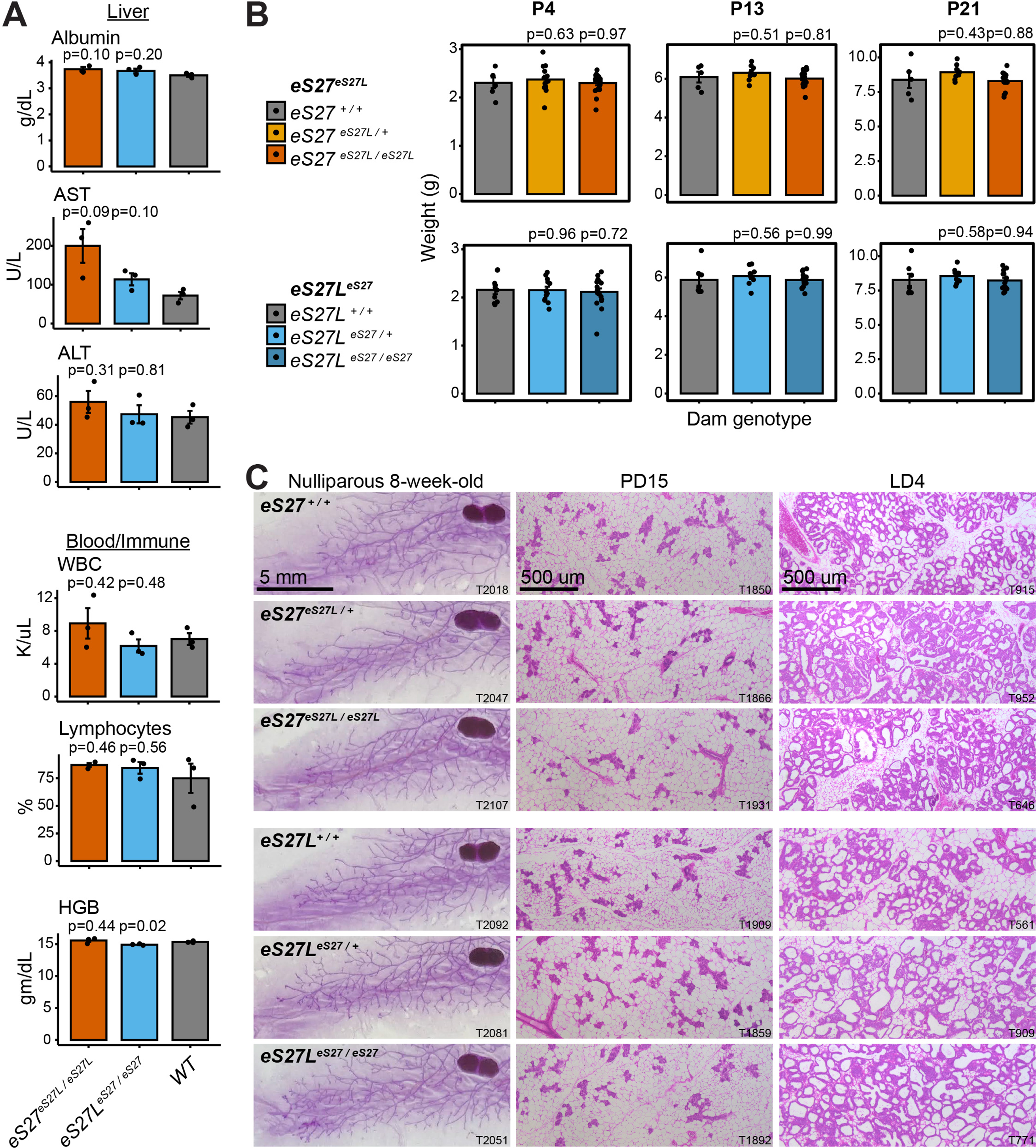
Homogenization of *eS27* and *eS27L* does not impact tissues that preferentially express one paralog. **(A)** Selected biomarkers from bloodwork performed on 10-16-week-old homozygous males and age-matched wild-type controls from the *eS27^eS27L^* and *eS27L^eS27^* mouse lines. See Supplementary Table 3 for additional biomarkers. AST = aspartate aminotransferase, ALT = alanine aminotransferase, WBC = white blood cells, HGB = hemoglobin. n = 3 biological replicates (individual animals) per genotype. Significance relative to WT was assessed by t-test. **(B)** Mean pup weight per litter at postnatal day (PD) 4, 13, and 21, grouped by dam genotype. Only pups from a dam’s first litter are included. n = 5-19 litters per dam genotype and timepoint. Significance relative to WT was assessed by t-test. **(C)** Representative images of carmine alum-stained or hematoxylin-eosin (H&E)- stained mammary glands at the indicated stages. T = animal ear tag number. See also Figure 6-- Source Data 1-18.

We considered the possibility that female mice housed under standard laboratory conditions may have a substantial reserve of lactation capacity, which could mask a moderate effect of *eS27(L)* homogenization on mammary gland development and function. We therefore used carmine alum staining and hematoxylin-eosin staining to assess the gross morphology and mammary fat pad filling of nulliparous, pregnant, and lactating females (Figure 6C). No anomalies in mammary gland morphogenesis were detected, with all genotypes exhibiting similar epithelial branch length and number, fat pad filling, alveolar size, and alveolar wall thickness. From this evidence, we conclude that homogenization of either the *eS27* or *eS27L* locus has no effect on overall mouse fitness and also no effect on the morphology or function of tissues that preferentially express either *eS27* or *eS27L*. These findings suggest that the eS27 and eS27L proteins are functionally similar in the setting of normal organism physiology, even in tissues that preferentially express one paralog.

### *eS27* and *eS27L* homogenization do not affect health in later life or in response to genotoxic stress

Having assessed for phenotypes in young homogenized male and female mice, we considered the possibility that the eS27 and eS27L proteins may function similarly under optimal conditions of homeostasis in young animals, but could act differently to confer an evolutionary benefit at a later age or under stress. To assess this hypothesis, we co-housed *eS27^eS27L^* and *eS27L^eS27^* homozygous, heterozygous, and wild-type male littermates under standard husbandry conditions (see Methods) until 9-10 months of age. At that time, we weighed the mice and performed a complete blood count and liver panel, again targeting the organ systems with preferential *eS27* or *eS27L* expression. No clinically significant differences were observed between genotypes in any of the included assays (Figure 7A, Supplementary Table 6). Thus, homogenization of *eS27* or *eS27L* has no detectable effect on the physiology of these organs, even later in life. These findings diminish the likelihood that there might be functional differences between the *eS27* and *eS27L* protein sequences that have cumulative effects over the mouse lifespan.

**Figure 7:**
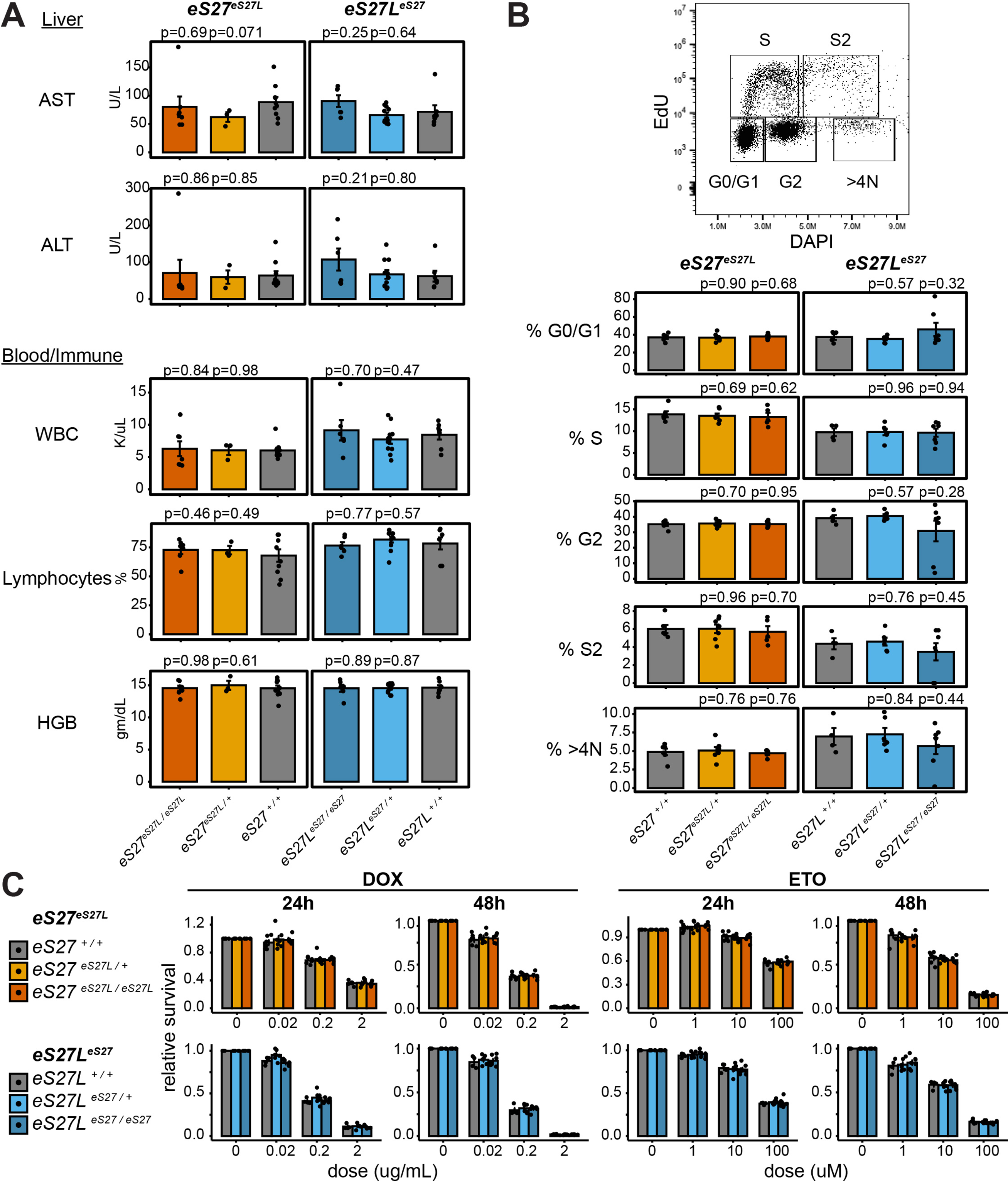
*eS27* and *eS27L* homogenization does not affect physiology at later age or impact response to genotoxic stress. **(A)** Biomarkers from 9-10-month-old *eS27^eS27L^* and *eS27L^eS27^* homozygous and heterozygous males and age-matched wild-type (WT) controls. n=3-11 animals per genotype. See Supplementary Table 5 for additional biomarkers. AST = aspartate aminotransferase, ALT = alanine aminotransferase, WBC = white blood cells, HGB = hemoglobin. **(B)** Percent of singlet mouse embryonic fibroblasts (MEFs) in each cell cycle phase as measured by EdU/DAPI flow cytometry. n=4-7 biological replicates (MEF lines isolated from single embry- os) **(C)** Fraction of viable MEFs relative to untreated controls after doxorubicin (DOX) or etoposide (ETO) treatment. n=4-7 biological replicates; n=2 technical replicates (separate wells) per biological replicate. For all panels, significance relative to WT was assessed by t-test. For (C), no comparisons were significant at p < 0.05.

Lastly, we tested the effects of stress stimuli on *ex vivo* cells derived from *eS27^eS27L^* and *eS27L^eS27^* mice. It has previously been reported that p53 differentially regulates *eS27* and *eS27L* expression (He & Sun, 2007; Li et al., 2007; Xiong et al., 2011). Furthermore, eS27 and eS27L proteins reportedly bind and regulate the p53-regulating ubiquitin ligase Mdm2 with different affinity *in vitro*, and thus may form distinct feedback loops impacting p53 activity (Xiong et al., 2011). Even though the expression patterns of *eS27* and *eS27L* that we observed are not likely driven by p53 (Figure 2F-G), we hypothesized that functional differences between the eS27 and eS27 proteins might be revealed under the types of genotoxic or cell cycle-related stress conditions for which p53 is classically a master regulator of response. We isolated mouse embryonic fibroblasts (MEFs) from homozygous, heterozygous, and wild-type mice of the *eS27^eS27L^* and *eS27L^eS27^* lineages. We first analyzed their distribution across cell cycle phases when cultured *in vitro*, and found that MEFs of all genotypes had similar frequency in each phase (Figure 7B). We then treated them with varying doses of doxorubicin and etoposide, two chemotherapeutic drugs that induce DNA damage and consequently activate *p53*. The number of viable and metabolically active cells was assessed after 24-48 hours of drug treatment (Figure 7C). Higher doses of doxorubicin and etoposide consistently resulted in decreased cell viability, yet the same degree of effect was seen across all genotypes. Thus, for the purposes of cellular survival and proliferation under doxorubicin or etoposide treatment, the eS27 and eS27L proteins also appear to be interchangeable.

## Discussion

In this work, we set out to test two categories of hypotheses regarding the evolutionary retention of a mammalian RP paralog pair: either that the paralogs encode functionally distinct proteins; or that their essentiality is due to dosage sharing or paralog buffering. We observed that *eS27* and *eS27L* mRNA abundance is inversely correlated and cell type-dependent across healthy mouse tissues, showed that eS27- and eS27L-ribosomes differentially associate with transcripts of cell cycle-related genes, and demonstrated that loss-of-function alleles of *eS27* and *eS27L* are both homozygous lethal but manifest at different developmental stages. Based on these findings, divergent protein functions and dosage sharing through subfunctionalized expression were both plausible reasons for *eS27* and *eS27L* conservation, while paralog buffering was not. Ultimately, after extensive examination of *eS27^eS27L^* and *eS27L^eS27^* mice, we concluded that the eS27 and eS27L proteins are functionally interchangeable. Together, these results suggest that *eS27* and *eS27L*, which possibly arose alongside other RP gene duplicates during a whole-genome duplication, have been evolutionarily retained because the divergence of their expression patterns has resulted in both genes becoming necessary for achieving adequate total expression of the RP across cell types (Figure 8).

**Figure 8:**
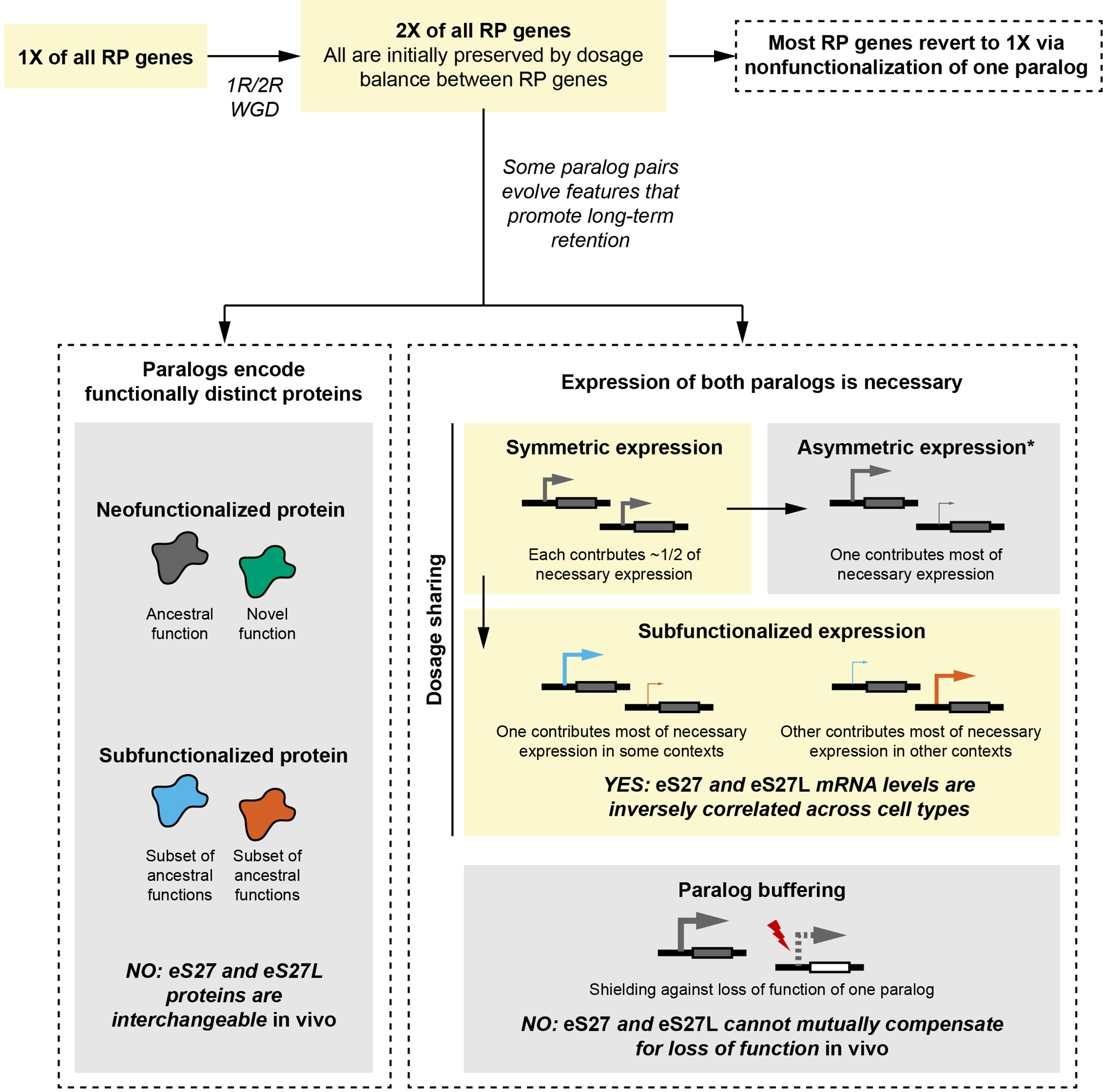
Comparison of empirical *eS27* and *eS27L* characteristics with hypothesized evolutionary trajectories. Yellow boxes indicate the most likely trajectory for *eS27* and *eS27L*. In brief, a whole-genome duplication (WGD) during early vertebrate evolution may have duplicated all RP genes. This mechanism is probable because the maintained dosage balance between RPs would have promoted initial preservation of duplicates. While most RP genes must then have reverted to single genes via nonfunctionalization of one paralog, some paralog pairs evolved features that promoted retention. Immediately after duplication, *eS27* and *eS27L* probably had similar regulatory elements and thus would have expressed symmetrically, but our findings suggest that eS27 and *eS27L* now exhibit subfunctionalized expression that renders both paralogs necessary to achieve the requisite total expression of this RP. We found no evidence of functional differences between eS27 and eS27L protein, nor successful compensation by either paralog for a loss-of-function of the other. Not pictured here are neofunctionalized expression and beneficial dosage increase; while these modes of paralog retention have been observed for other genes, they are less relevant for RP paralogs if it is assumed that excess dosage of an individual RP gene is not advantageous. *Symmetric expression frequently shifts towards asymmetric expression, which can be an intermediate state towards nonfunctionalization of the minor paralog.

Several questions arise when considering the *in vivo* homogenized mouse outcomes alongside the molecular findings reported here and in previous literature. We detected preferential association of eS27- or eS27L-ribosomes with cell cycle-related transcripts, yet replacing eS27 protein with eS27L and vice versa had no detectable impact on cell cycle progression (Figure 7B). One explanation is that eS27- and eS27L-ribosomes may have cell cycle-dependent abundance and therefore differentially encounter cyclically expressed transcripts, a consideration that remains to be addressed in future work. Another possibility is that eS27 and eS27L proteins do affect ribosome affinity for specific transcripts, but other pathways compensate to maintain a normal cell cycle in homogenized cells.

From previous literature, some evidence may suggest that eS27 and eS27L do have distinct protein functions, especially in p53-related signaling pathways and apoptotic processes resulting from genotoxic stimuli (He & Sun, 2007; Li et al., 2007; Xiong et al., 2018, 2020; Zhao et al., 2018).

Depletion or overexpression of *eS27L* were reported to impact p53 activity differently than *eS27* depletion or overexpression (Xiong et al., 2011, 2014). Purified eS27 and eS27L protein bind the p53 ubiquitin ligase Mdm2 *in vitro* with different affinity and are degraded by Mdm2 at different rates, which depends on the N-terminal portion of eS27 and eS27L where the differing residues reside (Xiong et al., 2011). Also, knockdown of *eS27L* but not *eS27* was reported to reduce the levels of DNA repair proteins FANCD2 and FANCI (Sun et al., 2020). However, in our comparison of WT and homogenized cells under genotoxic stimuli known to activate the *p53-Mdm2* axis (Figure 7C), we did not detect any differences in cell survival that would suggest functional divergence between the eS27 and eS27L proteins. We acknowledge the possibility that the two proteins could have distinct functions under environmental conditions not tested here, or that the effects of homogenization are masked by compensation in other cellular pathways. Nevertheless, the normal physiology of both homogenized mouse lines until at least 9-10 months of age suggests that total expression of *eS27* and *eS27L* impacts *in vivo* fitness more so than differences in protein characteristics. Importantly, our findings regarding homogenization involved no knockdown, knockout, or overexpression of RPs, thereby minimizing any indirect effects of perturbing RP expression.

The conclusion that *eS27* and *eS27L* likely persisted across evolution due to dosage sharing emphasizes the principle that distinct protein sequences and distinct expression patterns do not always indicate a paralog with tissue-specific protein functions. With the greater ease of genetic editing afforded by CRISPR, it would be fascinating to apply the endogenous epitope tagging and homogenization approaches used here to other mammalian RP paralogs that also have tissue- or cell type-dependent expression. *uL3L* (*Rpl3l*), for example, is an RP paralog that only expresses in skeletal and cardiac myocytes, where *uL3* (*Rpl3*) is not expressed (Guimaraes & Zavolan, 2016). While knockdowns and knockouts of *uL3L* in cells and mice have been characterized (Chaillou et al., 2016; Kao et al., 2021; Milenkovic et al., 2021), a homogenized *uL3L^uL3^* mouse allele would be invaluable for determining whether uL3L-ribosomes have myocyte-specific functions. As another example, *uL16L* (*Rpl10l*) and *eL39L* (*Rpl39l*) are mainly expressed in the testes and likely compensate during spermatogenesis for their respective progenitor genes, *uL16* (*Rpl10*) and *eL39* (*Rpl39*); the latter are encoded on the X chromosome and are therefore not expressed after meiotic X chromosome inactivation (Guimaraes & Zavolan, 2016; Sugihara et al., 2010). Knockouts of each have been characterized *in vivo* and *in vitro*, respectively, and exogenous expression of *uL16* partially rescues spermatogenesis in the *uL16L* knockout (Jiang et al., 2017; Zou & Qi, 2021; Zou et al., 2021). A homogenization approach would aid in determining whether the residual spermatogenesis defect is because the rescue does not precisely recapitulate the endogenous expression pattern of *uL16L*, or because *uL16* lacks some spermatogenesis-specific function of the *uL16L* protein. It is important to note that the properties of *eS27* and *eS27L* observed here do not necessarily extend to all mammalian RP paralogs, and that some mammalian RP paralog pairs, such as *uL3/uL3L* and *eL22/eL22L* (*Rpl22*/*Rpl22l*), have more differences in amino acid sequence than *eS27/eS27L*. Our work reinforces the importance of directly comparing protein function between RP paralogs in an endogenous context.

Lastly, our findings raise questions about the two *eS27* copies that have long been asked about gene duplication in general: did the ancestral *eS27* have some characteristic that led to its duplication and preservation? Is the existence of two copies with divergent expression but equivalent proteins evolutionarily beneficial? Or did their preservation and cell type-dependent expression patterns result from genetic drift without adaptive selection? It should first be noted that gene duplication is common (Lynch & Conery, 2000), and that few duplicates persist while most degrade. It should also be noted that not all gene features arising during evolution are necessarily beneficial, and that neutral evolution may play a large role in determining which duplicates persist and how their regulatory elements evolve (Lynch, 2007). RPs in general may have a propensity for paralog retention: in *Paramecium* and yeast, for example, duplicates of highly translated genes or components of protein complexes often persist, including ribosome, histone, or cytoskeleton proteins (Aury et al., 2006; Wapinski et al., 2007). Among present-day mammalian RPs, however, existing as a paralog pair puts *eS27* in the minority. A final curious observation is that, in vertebrate fish species that likely underwent additional WGDs, four or even eight eS27 copies have been reported whereas most other RPs have reverted to fewer copies (Kuang et al., 2020; Manchado et al., 2007). Is there something about eS27 that causes its paralogs to persist when other RP paralogs generated by WGD do not? Perhaps it is advantageous to have multiple copies whose expression could be more finely regulated (Greer et al., 2000), or perhaps the ancestral eS27 protein was already pleiotropic in its expression or protein function and was thus amenable to divergence into two genes with different expression and possibly subtle protein differences (Conant & Wolfe, 2008; Prince & Pickett, 2002). More work is needed to further probe the evolution and function of mammalian RP paralogs, and to provide additional comparisons between theoretical and experimental perspectives on paralog evolution. As a paradigm, the studies we report here set the groundwork for future investigation of RP paralog function.

### Data and Material Availability

All plasmids, cell lines, and mouse lines generated in this study are available upon request addressed to the corresponding author. All other reagents and materials used in this study are commercially available, and supplier information has been provided in the Methods section. Ribosome profiling sequencing data have been deposited in GEO under accession code GSE201845. Other data generated in this study are provided in the supplementary materials and Source Data files. Code used for data analysis is available at https://gitfront.io/r/adelefxu/f94QE89EJwyp/eS27-paralogs/.

## Materials & Methods

### Animal husbandry

All animal work was reviewed and approved by the Stanford Administrative Panel on Laboratory Animal Care (APLAC, protocol #27463). The Stanford APLAC is accredited by the American Association for the Accreditation of Laboratory Animal Care. All mice used in the study were housed at Stanford University except where otherwise noted. CRISPR-edited mouse lines were generated at the Gladstone Institute Transgenic Gene Targeting Core (San Francisco, CA). All animal procedures were approved by the Institutional Animal Care and Use Committee at the University of California, San Francisco (protocol #AN180952-01B). Mouse lines were maintained on a C57BL6/J background unless otherwise stated. *Trp53^null^* (Jacks et al., 1994) mice were purchased from Jackson Laboratories (strain #002101, Bar Harbor, ME). Mice were housed under 12-hour light-dark cycles with ad libitum irradiated chow (Teklad 2018SX, Envigo, Madison, WI), acidified water, and filtered air flow. For timed pregnancies, 1-2 female mice were housed overnight with 1 adult male and examined daily for vaginal plugs. Embryo stage was considered to be E0.5 on the day the vaginal plug was observed. Mice used in the same experiment were colony-matched, and also littermate-matched whenever possible. Mice were euthanized by CO2 inhalation and confirmed by cervical dislocation per APLAC guidelines.

Genotyping was performed using standard PCR protocols for MyTaq HotStart Red Mix (Bioline BIO- 25048, Memphis, TN) with primers listed in Supplementary Table 7.

### Reanalysis of RNA-seq datasets

Analysis scripts are available on GitHub (see “Data and Material Availability”). Gene count matrices and cell or sample type annotations were downloaded from the Mouse Cell Atlas (Han et al., 2018), Tabula Muris (Tabula Muris Consortium et al., 2018), Bach et al (Bach et al., 2017), and Fu et al (Fu et al., 2015). For scRNA-seq datasets, read counts from single cells were pooled. Total reads of each RP gene in a cell type were normalized by the total read count of all RP genes in the cell type to normalize for the cell type-specific rate of ribosome production, then multiplied by 100. For non-RP genes, normalized read counts are reported as reads per million.

### RT-qPCR from mouse mammary glands, heart, liver, and brain

For Figure 2A, whole mammary gland tissue was harvested from abdominal glands at the indicated timepoints. The central lymph node was removed, and total RNA was isolated using NucleoSpin RNA (Macherey-Nagel 740955.50, Düren, Germany) according to the manufacturer’s instructions. For Figure 2E-F, tissues were harvested, cut into ∼3 mm pieces, and snap-frozen in liquid nitrogen. For mammary glands, the central lymph node was removed prior to freezing. Snap-frozen tissues were powderized in a ceramic mortar and pestle while submerged in liquid nitrogen. Powder was suspended in TRIzol (ThermoFisher 15596-018, Waltham, MA). RNA was extracted according to manufacturer protocol and isolated using the PureLink RNA Mini kit (ThermoFisher 12183018A).

Samples were treated with Turbo DNAse and inactivated (ThermoFisher AM1907).

For all samples, 1 ug of RNA was reverse-transcribed using iScript RT Supermix (Bio-Rad 1708841, Hercules, CA). qPCR was performed using SsoAdvanced SYBR Green Super Mix (Bio-Rad 1725270) on a Bio-Rad CFX384 using the primers listed in Supplementary Table 7. Two technical replicates were performed for each of the three biological replicates per condition. Ct values were normalized to a housekeeping gene or another RP gene as stated in figure legends, and displayed as a fold difference relative to a reference sample.

### Mouse embryonic stem cell (mESC) culture

E14Tg2a.4 mESCs (Smith and Hooper, 1987) were a gift from Thom Saunder’s lab (University of Michigan). Cells were cultured in Knockout DMEM (ThermoFisher 10829-018) supplemented to a final concentration of 15% ES-qualified fetal bovine serum (MilliporeSigma ES-009-B, Burlington, MA), 1% non-essential amino acids (MilliporeSigma TMS-001-C), 2 mM L-glutamine (MilliporeSigma TMS- 002-C), 1% penicillin/streptomycin (ThermoFisher 15140-122), 55 uM beta-mercaptoethanol (ThermoFisher 21985-023), and 1000 U/mL mouse leukemia inhibitory factor (mLIF, Gemini 400-495 10^7, West Sacramento, CA). mESCs were not routinely tested for mycoplasma contamination. Cell line identity was confirmed daily by observing colony morphology. Media was changed daily and cells were passaged every two days. To passage, plates with ∼70% confluent colonies of mESCs were washed in Dulbecco′s Phosphate Buffered Saline (DPBS) (ThermoFisher 14190-250) and trypsinized (0.05% trypsin, dilution of ThermoFisher 15400-054 in DPBS) for 5 minutes at 37 C. Trypsin was neutralized by a double volume of media. Cells were immediately dissociated by vigorous pipetting, pelleted at 200 g for 3 minutes at room temperature, and resuspended in fresh warmed media for plating. Fresh plates were pre-coated at 37 C overnight with 0.1% gelatin (MilliporeSigma ES-006-B), which was aspirated prior to plating cells. Unless otherwise stated, cells were plated at a density equivalent to 5 x 10^6^ cells per 10 cm plate.

### Density gradient fractionation

Gradient lysis buffer: 20 mM Tris pH 7.5 (ThermoFisher AM9850G, AM9855G), 150 mM NaCl (ThermoFisher AM9760G), 15 mM MgCl2 (ThermoFisher AM9530G), 1% v/v Triton-X 100 (MilliporeSigma X100-500ML), 8% v/v glycerol (MilliporeSigma G6279), 1 mM DTT (MilliporeSigma 43815), 1X Complete mini protease inhibitor EDTA-free (Roche 11836170001, Basel, Switzerland), 0.5% w/v deoxycholate (MilliporeSigma S1827), 100 ug/mL cycloheximide (MilliporeSigma C7698), 0.02 U/uL Turbo DNAse (ThermoFisher AM2239), 0.2 U/uL Superase RNAse Inhibitor (ThermoFisher AM2696) in Ultrapure distilled water (ThermoFisher 10977-015).

Gradient sucrose buffer: 10 or 45% w/v sucrose (Fisher 8510500GM), 20 mM Tris pH 7.5 (ThermoFisher AM9850G, AM9855G), 100 mM NaCl (ThermoFisher AM9760G), 15 mM MgCl2 (ThermoFisher AM9530G), 1 mM DTT (MilliporeSigma 43815), 100 ug/mL cycloheximide (MilliporeSigma C7698) in Ultrapure distilled water (ThermoFisher 10977-015).

A 10 cm plate of mESCs at ∼50% confluence was treated with 100 ug cycloheximide (CHX) for 2 minutes at 37 C. Cells were washed, trypsinized, and neutralized with media as described above, except that all buffers contained 100 ug/mL CHX. The cells were pelleted at 200 g x 3 min at 4 C, washed with ice-cold DPBS+CHX, and pelleted again. To the pellet, 400 uL gradient lysis buffer (above) was added and vortexed for 30 seconds with 30 seconds rest on ice for 3 cycles, then incubated for 30 minutes rotating at 4C. The lysate was clarified by centrifuging at 700 g for 5 min at 4C, then again at 7000 g for 5 min at 4 C. RNA concentration was measured by Nanodrop 2000 (ThermoFisher) and samples were adjusted to equal concentrations using additional lysis buffer. 250 uL of clarified lysate was layered onto a 10-45% sucrose gradient (see above), which was made on a Biocomp Model 108 Gradient Master (Colorado Springs, CO). Gradients were spun on a Beckman SW- 41 rotor (Indianapolis, IN) at 40,000 rpm for 2.5 hours at 4 C. After centrifugation, gradients were fractionated using a density gradient fraction system (Brandel BR-188, Gaithersburg, MD) with measurement of UV absorbance. Fractions were precipitated using the ProteoExtract protein precipitation kit (MilliporeSigma 539180) per manufacturer protocol and redissolved in 2X Laemmli buffer (Fisher 50-196-784). For Western blot, an equal volume of each fraction was loaded onto an SDS-PAGE gel.

### Western blot

Western blot lysis buffer: 25 mM Tris pH 7.5 (ThermoFisher AM9850G, AM9855G), 150 mM NaCl (ThermoFisher AM9760G), 15 mM MgCl2 (ThermoFisher AM9530G), 1% v/v Triton-X 100 (MilliporeSigma X100-500ML), 8% v/v glycerol (MilliporeSigma G6279), 1 mM DTT (MilliporeSigma 43815), 1X Complete mini protease inhibitor EDTA-free (Roche 11836170001), 0.5% w/v deoxycholate (MilliporeSigma S1827), 0.02 U/uL Turbo DNAse (ThermoFisher AM2239), 0.2 U/uL Superase RNAse Inhibitor (ThermoFisher AM2696) in Ultrapure distilled water (ThermoFisher 10977-015).

Unless otherwise stated, cells were washed, trypsinized, and neutralized with media as described above. The cells were pelleted at 200 g for 3 min at 4C, washed with ice-cold DPBS, pelleted again, and washed again in DPBS. Western blot lysis buffer (above) was added. Cells and lysis buffer were vortexed for 30 seconds and rested on ice for 30 seconds for three cycles, then incubated at 4 C for 15 minutes. Lysates were clarified by centrifuging at 7000 g for 5 minutes at 4C. Unless otherwise stated, total protein in each sample was quantified by bicinchoninic acid assay (ThermoFisher 23225) per manufacturer protocol and samples were normalized to equal total protein.

Samples were resolved on a 4%–20% Tris-glycine gradient SDS-PAGE gel (Bio-Rad 5671095) and transferred to a polyvinylidene difluoride membrane (Bio-Rad 1704273). Membranes were blocked for 1 hour at room temperature with 5% milk in phosphate-buffered saline (PBS) (Fisher BP2944100) with 0.1% Tween-20 (MilliporeSigma P9416) (PBST). Blots were incubated for 16 h at 4°C with the following primary antibodies at 1:1000 dilution in 5% BSA/PBST, unless stated otherwise: anti-eS27 (1:10000, ThermoFisher PA5-18092), anti-eS27L (ProteinTech 15871-1-AP, Rosemont, IL), anti-B-actin (Cell Signaling 3700S, Danvers, MA), anti-uS3 (Abcam ab77330, Cambridge, United Kingdom), anti- FLAG (MilliporeSigma F3165), anti-GAPDH (1:5000, ThermoFisher AM4300), anti-p53 (Leica Biosystems CM5, Wetzlar, Germany). Membranes were washed 3 times for 10 min in PBST before incubation for 30 min at room temperature with secondary antibodies coupled to horseradish peroxidase at 1:10000 dilution in 5% milk/PBST: donkey anti-mouse (GE Healthcare NA931-1ML, Chicago, IL), donkey anti-rabbit (GE Healthcare NA934-1ML), chicken anti-goat (R&D Systems HAF019, Minneapolis, MN). Membranes were washed 3 times for 10 min in PBST before detection using Clarity Western ECL Substrate (Bio-Rad 170-5061) and imaging on a ChemiDoc MP (Bio-Rad 17001402).

### Generation of CRISPR-edited mESCs

The CRISPR strategy for inserting 3xFLAG C-terminally at the endogenous eS27 or eS27L locus in mESCs via exon replacement was designed as follows: first, two guide RNA (gRNA) recognition sites were identified that flanked the exon of eS27 or eS27L containing the stop codon, using on- and off-target gRNA site scoring algorithms (Doench et al., 2016; Hsu et al., 2013) implemented in Benchling (San Francisco, CA). gRNA sites were only used if they had no other high-probability predicted cut sites throughout the mouse genome. Each 20 nt gRNA sequence was each cloned into a PX459 (Ran et al., 2013) (Addgene 62988, Watertown, MA) backbone digested with BbsI (Thermo FD1014), with a single upstream G nucleotide preceding the gRNA sequence since this has been shown to improve cutting efficiency (Ran et al., 2013). To construct the homology-directed repair template, the sequence between the two gRNA cut sites was cloned along with 300 bp homology arms on each end. Immediately preceding the stop codon in the repair template, a sequence was inserted to encode a 2xGGGS linker and a 3xFLAG peptide. The gRNA recognition sites or protospacer adjacent motifs (PAM) on the repair template were modified at silent coding positions or non-coding positions with low evolutionary conservation to prevent cutting of the repair template or of the repaired genomic DNA. To deliver the repair template, an unmodified PAM and gRNA recognition sequence were appended to each end of the repair template, distal to the homology arms. Distal to the appended gRNA and PAM sequences on each side, 10 additional bases were appended to buffer against small deletions that occur with TOPO cloning. This construct was then inserted into a non-expressing pCR4Blunt-TOPO backbone (ThermoFisher 450031) such that the Cas9-gRNAs expressed from the PX459 plasmids would cleave the repair template as a linear dsDNA from the circular pCR4Blunt- TOPO plasmid (J.-P. Zhang et al., 2017).

At passage number 28, 10^6^ mESCs were transfected with the two PX459-based plasmids harboring a Cas9-puromycin fusion construct and gRNAs flanking the targeted exon (0.5 ug each), and one pCR4Blunt-TOPO-based plasmid harboring the linearizable repair template (2 ug) (see Supplementary Table 7 for sequences). The three combined plasmids were diluted in 100 uL Opti-MEM Reduced Serum Medium (ThermoFisher 11058021). In parallel, 7.5 uL of Lipofectamine 2000 (ThermoFisher 11668-019) was diluted in 100 uL Opti-MEM. The plasmid/Opti-MEM and Lipofectamine/Opti-MEM were combined and incubated for 20 minutes at room temperature. Cells were trypsinized as described above, resuspended in 250 uL Opti-MEM, added to the plasmid-Lipofectamine complexes for 10 minutes at room temperature, and plated into one 12-well. Media was changed after 4 hours. At 24 hours after transfection, cells were treated with media containing 1 ug/mL puromycin (Millipore P8833). At 48 hours after transfection, fresh media containing puromycin was changed in. At 72 hours after transfection, puromycin-free media was changed in. At 96 hours after transfection, cells were washed, trypsinized, dissociated, and plated at 1000 cells per 10 cm plate to form colonies derived from single cells. At 7 days after sparse plating, individual colonies were lifted using a pipet tip, dissociated at 37 C in 0.025% trypsin-EDTA in DPBS, replica plated in gelatinized 96-well plates in regular mESC media, and screened by genomic DNA PCR and Western blot for desired edits.

Importantly, PCR primers were designed such that at least one primer bound distal to the 300 bp homology arms, to avoid amplifying residual repair template.

### FLAG-IP ribosome profiling

IP lysis buffer: 25 mM Tris pH 7.5 (ThermoFisher AM9850G, AM9855G), 150 mM NaCl (ThermoFisher AM9760G), 15 mM MgCl2 (ThermoFisher AM9530G), 1% v/v Triton-X 100 (MilliporeSigma X100-500ML), 8% v/v glycerol (MilliporeSigma G6279), 1 mM DTT (MilliporeSigma 43815), 1X Complete Mini Protease Inhibitor EDTA-free (Roche 11836170001), 0.5% w/v deoxycholate (MilliporeSigma S1827), 200 ug/mL cycloheximide (MilliporeSigma C7698), 0.02 U/uL Turbo DNAse (ThermoFisher AM2239), 0.2 U/uL Superase RNAse Inhibitor (ThermoFisher AM2696) in Ultrapure distilled water (ThermoFisher 10977-015).

Sucrose cushion buffer: 25 mM Tris pH 7.5 (ThermoFisher AM9850G, AM9855G), 150 mM NaCl (ThermoFisher AM9760G), 15 mM MgCl2 (ThermoFisher AM9530G), 1 mM DTT (MilliporeSigma 43815), 1X Complete Mini Protease Inhibitor EDTA-free (Roche 11836170001), 200 ug/mL cycloheximide (MilliporeSigma C7698) in Ultrapure distilled water (ThermoFisher 10977-015).

Wash buffer 1: equivalent to IP lysis buffer but omitting glycerol, protease inhibitor, RNAse inhibitor, and DNAse

Wash buffer 2: equivalent to wash buffer 1, but with 300 mM NaCl

Western elution buffer: 4% v/v SDS (MilliporeSigma 436143), 125 mM Tris pH 7.0 (ThermoFisher AM9850G) in Ultrapure distilled water (ThermoFisher 10977-015).

CRISPR-edited mESCs and control lines were grown to ∼50% confluence in 2x15 cm plates.

Three biological replicates (clones) were included for each genotype. Cells were treated with 100 ug/mL cycloheximide (CHX) for 2 minutes at 37 C, then washed twice with 20 mL DPBS containing 100 ug/mL CHX. Cells were scraped into 5 mL DPBS+CHX and centrifuged at 200 g for 3 minutes at 4C, then washed again in DPBS+CHX. 800 uL IP lysis buffer without Superase RNAse Inhibitor was added to the pellet and vortexed for 30 seconds with 30 seconds rest for 3 cycles, then rotated at 4 C for 30 minutes. Lysate was clarified by centrifuging at 700 g for 5 min at 4C, then again at 7000 g for 5 min at 4C. RNA concentration was measured by Nanodrop 2000 (Thermo). By adding more IP lysis buffer without RNAse inhibitor, the clarified lysate was adjusted to a concentration of 1500 ng/uL in 600 uL total volume. 6 uL RNase A (Thermo EN0531) and 3.6 uL RNAse T1 (Thermo EN0541) were added, and the digestion reaction was rotated for 30 minutes at room temperature. RNAse digestion was quenched by placing it on ice and adding 30 uL of Superase RNAse Inhibitor. Two 250 uL aliquots of digested lysate were each layered over 750 uL of sucrose cushion buffer (see above). The cushions were ultracentrifuged in a TLA 120.2 rotor (Beckman, Indianapolis, IN) at 100,000 rpm for 1 hour at 4 C. Each pellet was rinsed once with 1 mL ice-cold Ultrapure water, then resuspended by pipetting and shaking in 250 uL IP lysis buffer containing Superase RNAse Inhibitor. An aliquot of the resuspended pellet was reserved in TRIzol (ThermoFisher 15596-018) as the total RPF fraction. The two resuspended pellets from each replicate were combined and added to 200 uL mouse IgG beads (MilliporeSigma A0919) which had been equilibrated twice for 5 min each at 4 C in an equal volume of IP lysis buffer with RNAse inhibitor. The samples were pre-cleared with the IgG beads for 1 hour at 4 C on a rotator, then transferred to 200 uL anti-FLAG beads (MilliporeSigma A2220), which had been equilibrated twice for 5 min each at 4 C in an equal volume of IP lysis buffer with RNAse inhibitor. The immunoprecipitation reaction was rotated for 2 hours at 4C, washed with 400 uL wash buffer 1 three times for 5 min each at 4C, and washed with 400 uL wash buffer 2 three times for 5 min each at 4 C. For ribosome profiling, an aliquot of the beads was incubated in TRIzol for 5 minutes at room temperature and reserved as the IP RPF fraction. For Western blot, an aliquot of the beads was heated with shaking to 95 C in Western elution buffer (see above) for 5 min.

For Western blot, an aliquot of the cushion supernatant and eluate were precipitated using the ProteoExtract protein precipitation kit (Calbiochem 539180) per manufacturer protocol and redissolved in 2X Laemmli buffer (Fisher 50-196-784). 1% of the sample volume was loaded for lysate, supernatant, pellet, and flow-through samples. 10% of the sample volume was loaded for eluate samples.

Ribosome profiling libraries were prepared following the published protocol of McGlincy and Ingolia (McGlincy & Ingolia, 2017) with modifications as stated below, using oligonucleotides synthesized by Integrated DNA Technologies (Coralville, IA) (Supplementary Table 7). Total RPFs and IP RPFs were extracted from TRIzol using the Direct-Zol Microprep Kit (Zymo R2060, Irvine, CA) according to the manufacturer protocol, omitting the DNAse digestion. RPF volume was adjusted to 90 uL with Ultrapure water. 10 uL 3M NaOAc pH 5.5 (ThermoFisher AM9740) and 2 uL of 15 mg/mL GlycoBlue (ThermoFisher AM9515) were added before precipitating the RPFs in 150 uL 100% isopropanol overnight at -80 C. Precipitated RPFs were pelleted at 21,000 g for 30 minutes at 4 C, washed with ice-cold 80% ethanol in water, dried at room temperature for 10 minutes, and dissolved in Ultrapure water. RPFs were denatured at 80 C for 90 sec in denaturing sample loading buffer (McGlincy & Ingolia, 2017), then incubated on ice for 5 min before running on a 15% Tris-borate-EDTA- urea (TBE-urea) polyacrylamide gel. Fragments were size-selected using NI-800 and NI-801 (McGlincy & Ingolia, 2017) as 26-34 nt markers. Gel slices were freeze-thawed for 30 minutes at -80 C, crushed, and extracted at room temperature overnight in 400 uL RNA extraction buffer (McGlincy & Ingolia, 2017), then re-extracted with an additional 200 uL RNA extraction buffer. The combined 600 uL extraction was precipitated with 2 uL GlycoBlue and 750 uL 100% isopropanol overnight at -80 C. The RPFs were pelleted, washed, and dried as described above, then dissolved in 4 uL of 10 mM Tris pH 8, dephosphorylated, and ligated to barcoded linkers per the published protocol. Between replicates, barcodes were permuted among the samples. Unreacted linker was deadenylated and digested per the published protocol. The barcoded RPFs were pooled within each replicate and purified on a Zymo Oligo Clean & Concentrator column (Zymo D4060) according to manufacturer protocol. Pooled RPFs were diluted to 100 ng/uL and 1 ug was used as input for rRNA depletion using RiboZero Gold (part of Illumina 20020598, San Diego, CA) per manufacturer protocol. rRNA-depleted RPFs were purified on a Zymo Oligo Clean & Concentrator column, then reverse-transcribed per the published protocol.

Template RNA was degraded by alkaline hydrolysis per the published protocol. cDNA was purified on a Zymo Oligo Clean & Concentrator column, denatured in denaturing sample loading buffer, and size- selected on a 10% TBE-urea gel, as marked by NI-800 and NI-801 that had been ligated and reverse- transcribed in parallel with the samples. Gel slices were freeze-thawed for 30 minutes at -80 C, crushed, and extracted at room temperature overnight in 400 uL DNA extraction buffer (McGlincy & Ingolia, 2017), then re-extracted with an additional 200 uL DNA extraction buffer. The combined 600 uL extraction was precipitated with 2 uL GlycoBlue and 750 uL 100% isopropanol overnight at -80 C. The cDNA was pelleted, washed, and dried as described above, then resuspended in 15 uL 10 mM Tris pH 8. cDNA was circularized by adding 2 uL 10X CircLigase I buffer, 1 uL of 1 mM ATP, 1 uL of 50 mM MnCl2, and 1 uL of CircLigase I (Lucigen CL4111K, Middleton, WI) to the 15 uL of cDNA and incubating at 60 C for 12 hours, then 80 C for 10 minutes. Circularized cDNA was purified on a RNA Clean & Concentrator column. cDNA concentration was quantified by qPCR per the published protocol except using SsoAdvanced Universal SYBR Green qPCR master mix (Biorad 1725274) and following manufacturer protocol. Library construction to add indexing primers was performed per the published protocol, using a different reverse primer for each replicate. PCR products were purified using a Zymo DNA Clean & Concentrator column (Zymo D4003). Size selection was performed on a 8% TBE-urea gel, with the lower bound marked by NI-803 (McGlincy & Ingolia, 2017) that had undergone library construction in parallel with the samples, and the upper bound at 170 nt as marked by O’Range 20 bp DNA ladder (Thermo SM1323). Gel slices were extracted as described above. DNA was precipitated as described above, except using 1.25 uL of 20 ug/mL glycogen (Themo 10814-010) instead of GlycoBlue in the precipitation and incubating at -80 C for 2 hours. The pellet was resuspended in 10 mM Tris pH 8. Library quality and concentration was analyzed on an Agilent 2100 Bioanalyzer (High-Sensitivity DNA) at the Stanford Protein and Nucleic Acid Facility. Libraries were sequenced by Novogene (Sacramento, CA) on an Illumina HiSeq 4000 with paired-end 150 bp reads.

### Ribosome profiling analysis

Analysis scripts are available on GitHub (see “Data and Material Availability”). Due to the short insert length, only analysis of Read 1 was necessary. cutadapt version 2.4 (Martin, 2011) was used to trim 3’ adapter sequences from Read 1 with parameters “-j 0 -u 3 -a AGATCGGAAGAGCACAGTCTGAACTCCAGTCAC --discard-untrimmed -m 15”. In-line barcodes were demultiplexed using fastx_barcode_splitter.pl (http://hannonlab.cshl.edu/fastx_toolkit/) with parameters “--eol”. Unique molecular identifiers and in-line barcodes were extracted using umi_tools version 1.0.1 (Smith et al., 2017) with parameters “extract --extract-method=string --bc-pattern=NNNNNCCCCC -- 3prime”. Reads were filtered by quality using fastq_quality_filter (http://hannonlab.cshl.edu/fastx_toolkit/) with parameters “-Q33 -q 20 -p 70 -z”. To remove reads originating from rRNA, transfer RNA (tRNA), and small nuclear RNA (snRNA), reads aligning to these sequences using bowtie2 version 2.3.4.3 (Langmead & Salzberg, 2012) with parameters “-L 18” were discarded. Remaining reads were aligned using bowtie2 with parameters “--norc -L 18” to a reference GRCm38/mm10 mouse transcriptome that was derived from UCSC/GENCODE VM20 knownCanonical annotations filtered for transcripts associated with at least one of the following: a Uniprot ID, a RefSeq ID, or an Entrez ID. PCR duplicates were removed using umi_tools. RPFs were parsed for uniquely aligned reads and grouped by read length. Ribosome A site positions were determined by offsetting the distance of the 5’ end of each read to canonical start sites in each length group and adding 4 nucleotides. RPF reads aligning to the coding sequence (CDS) of a transcript (excluding the first 15 codons and last 5 codons of each CDS) were counted, using the above transcriptome annotation.

Because genes encoded by mitochondrial DNA are translated by mitoribosomes in the mitochondrial lumen that are distinct from cytoplasmic ribosomes, reads mapping to these genes were excluded from further analysis. Transcripts with counts per million (CPM) > 2 were retained for downstream analysis. IP RPF and total RPF libraries were normalized separately by the trimmed mean of M-values method in edgeR (Robinson et al., 2010).

Differential RPF abundance and enrichment were analyzed using voom (Law et al., 2014) and limma (Ritchie et al., 2015). The following terminology describes the variables included in the analysis: “cell line” distinguishes eS27-FLAG, eS27L-FLAG, and WT mESCs; “biological replicate” distinguishes each mESC clone (three independently selected clones per cell line); and “fraction” distinguishes IP and total RPFs.

To identify transcripts that were differentially abundant between the total RPFs in eS27-FLAG, eS27L-FLAG, and WT mESCs, the following design matrix was used: “∼0 + cell line.” Contrast matrices were constructed for each pairwise comparison: eS27-FLAG versus WT, eS27L-FLAG versus WT, and eS27L-FLAG versus eS27-FLAG.

To identify transcripts that were differentially abundant between the IP and total RPFs of each cell line, the following design matrix was used: “∼0 + fraction.” Contrast matrices were constructed for each pairwise comparison: eS27-FLAG IP versus total, eS27L-FLAG IP versus total.

To identify transcripts that were differentially enriched by anti-FLAG IP from among the total RPFs in eS27-FLAG versus eS27L-FLAG mESCs, the following design matrix was constructed: “∼0 + cell line + cell line:biological replicate + cell line:fraction.” The following comparison was made: eS27L- FLAG:IP versus eS27-FLAG:IP.

For all comparisons, the data for the relevant samples were transformed using voom to remove mean-variance count heteroscedasticity. Using the respective design and contrast matrices described above, linear models were fitted to the data using limma with empirical Bayes moderation. Transcripts with significant differential RPF abundance or enrichment were defined as those with false discovery rate < 0.05 obtained using the Benjamini-Hochberg method.

Gene ontology enrichment analysis was performed using the goana function in limma after excluding a *Tmod3* transcript as a likely artifact of anti-FLAG IP (Supplementary Text, Supplementary Figure 3).

### Generation of genetically edited mouse lines

The CRISPR strategy to produce mice with eS27 or eS27L homogenization or truncation is analogous to the strategy described above for CRISPR insertion of 3xFLAG into mESCs. Highly specific gRNA recognition sites were selected that flanked exon 2 of either gene, which encodes the three residues that differ between the paralogs. Instead of a linearizable dsDNA template, an ssDNA repair template with 100 nt homology arms was synthesized by GeneWiz (South Plainfield, NJ). This repair template contained the modified codons to homogenize the targeted paralog. Additionally, the gRNA recognition sites or protospacer adjacent motifs (PAM) on the ssDNA repair template were modified at silent coding positions or non-coding positions with low evolutionary conservation to prevent cutting of the repaired genomic DNA. gRNA and ssDNA sequences are listed in Supplementary Table 7.

CRISPR-edited mouse lines were generated at the Gladstone Institute Transgenic Gene Targeting Core (San Francisco, CA). All animal procedures were approved by the Institutional Animal Care and Use Committee at the University of California, San Francisco (protocol #AN180952-01B). Superovulated female C57BL/6 mice (4 weeks old) were mated to C57BL/6 stud males. Fertilized zygotes were collected from oviducts and injected with Cas9 protein (20 ng/uL), two sgRNAs with recognition sites that flanked exon 2 (10 ng/uL each) (synthesized by IDT as Alt-R CRISPR-Cas9 crRNA, Coralville, IA), and one ssDNA repair template (10 ng/uL) into the pronucleus of fertilized zygotes. Injected zygotes were implanted into oviducts of pseudopregnant CD1 female mice. The targeted loci were PCR-amplified from genomic DNA of F0 animals, subcloned, and Sanger sequenced to identify successfully edited alleles. The *eS27^exon2del^* and *eS27L^exon2del^* alleles were recovered serendipitously from mice injected with reagents designed to produce the *eS27^eS27L^* and *eS27L^eS27^* alleles, which were also successfully recovered. F0 mice were backcrossed to wild-type C57BL6 males and females (Jackson Labs) for at least four generations.

### Adult mouse necropsy and bloodwork

For necropsy of 10-16-week-old adult male mice, mice were euthanized by CO2 asphyxiation and cardiac exsanguination. Whole cardiac blood was collected in EDTA-coated microtainers (Becton Dickinson 365974) and 1.5 mL plastic tubes for complete blood counts and serum biochemistry, respectively. Blood samples were analyzed as described below. Mice were routinely processed for gross examination. In addition to body weight, the following organs were weighed, and a percentage of body weight calculation was conducted for each organ: liver, spleen, heart, kidneys (left and right), testicles (left and right). Tissues were immersion-fixed in 10% neutral buffered formalin (Fisher Scientific) for 72 hours. Tissues containing bone were decalcified for 24 hours using Cal-Ex II Fixative/Decalcifier (Fisher CS511-1D). Formalin-fixed tissues were processed routinely, embedded in paraffin, sectioned at 5 µm, and stained with hematoxylin and eosin. The following organs were evaluated histologically by a board-certified veterinary pathologist blinded to the sample genotypes: liver, kidneys, heart, spleen, thymus, pancreas, salivary glands, lungs, thyroid, trachea, esophagus, tongue, haired skin (interscapular), testes, accessory sex glands (preputial, seminal vesicles, prostate), urinary bladder, brain, gastrointestinal tract, bone marrow (pelvic limb), nasal cavity, eyes, teeth, ears, vertebral column, spinal cord.

For bloodwork of aged adult mice, male mice were co-housed with male littermate controls until 9-10 months of age. Mice were euthanized by CO2 asphyxiation and cardiac exsanguination. Whole cardiac blood was collected as described above. Samples were handled at room temperature during collection and submitted for analysis within one hour of collection. Automated hematology was performed on a Sysmex XN-1000V hematology analyzer. Blood smears were made for all CBC samples, Wright-Giemsa stained, and reviewed by a clinical laboratory scientist. Manual differentials were performed as indicated by species and automated analysis. Liver panel analysis was performed on a Siemens Dimension EXL200/LOCI analyzer.

### Mammary carmine alum, H&E

After euthanasia, mammary glands were dissected from littermate-matched female mice at the indicated pregnancy and lactation stages. From each mouse, the left abdominal mammary gland was used for carmine alum staining as previously described (Plante et al., 2011). The right abdominal mammary gland was fixed overnight in 10% neutral-buffered formalin, washed in phosphate buffered saline with 0.2% w/v glycine, and stored in 70% ethanol until paraffin embedding, sectioning, and staining with hematoxylin-eosin.

### Mouse embryonic fibroblast isolation and culture

Mouse embryonic fibroblasts (MEFs) were isolated at E13.5 as previously described (Durkin et al., 2013) from heterozygous crosses of the eS27^eS27L^ and eS27L^eS27^ lines. MEFs were cultured in DMEM with high glucose (Gibco 11965), supplemented with 10% fetal bovine serum (MilliporeSigma TMS-013-B) and 1% penicillin/streptomycin (ThermoFisher 15140-122). MEFs were cultured at 37 C with 5% CO2 and passaged every 2-3 days following the same passaging protocol as performed for mESCs (above). MEFs were used for experiments at passage numbers 2-4.

### EdU/DAPI Flow Cytometry

MEFs were plated at 250,000 cells per T25 flask. After 72 hours, 5-ethynyl-2’-deoxyuridine (EdU) (Click Chemistry Tools 1381, Scottsdale, AZ) was added to a final concentration of 10 uM. The media was kept warm during the EdU addition and thoroughly mixed in the flask afterwards. The cells were incubated at 37 C for 2 hours, then washed once with 3 mL of 0.05% trypsin-EDTA in DPBS and trypsinized for 5 min at 37 C in 1 mL 0.05% trypsin-EDTA in DPBS. The trypsin was neutralized with 1 mL MEF media and pipetted thoroughly to obtain a single-cell suspension. The media and trypsin wash previously poured off from the flask was recombined with the cells. The cells were pelleted at 200 g for 3 minutes at room temperature and washed in 1 mL 1% bovine serum albumin (BSA) in DPBS. For each MEF line, 500K cells were fixed in 200 uL 4% paraformaldehyde in PBS (dilution of Fisher 43368- 9M in 1.33X PBS) for 15 minutes in the dark at room temperature. To the fixative reaction, 1 mL ice-cold 70% ethanol was added and incubated on ice for 10 minutes. Cells were washed twice in 1 mL DPBS + 0.1% Triton X-100 and resuspended in 50 uL DPBS + 0.1% Triton X-100. The click reaction master mix was prepared per manufacturer instructions (Click Chemistry Tools 1381) and 250 uL was added to each sample. The click reaction was incubated for 30 minutes in the dark at room temperature, then washed with 1 mL DPBS + 0.1% Triton X-100. The cells were resuspended in 150 uL DPBS + 0.1% Triton X-100 containing 4 ug/mL 4′,6-diamidino-2-phenylindole (DAPI) (Thermo 62248), filtered through a mesh-top tube, and incubated in the dark overnight at 4 C before running on a Novocyte Quanteon flow cytometer using NovoExpress 1.3.0 software (Agilent, Santa Clara, CA) at a flow rate of 14 uL/minute. Gating for singlets was performed in FlowJo (Ashland, OR) on forward scatter area (FSC-A) x forward scatter height (FSC-H), then DAPI area x height.

### Cell viability assays

MEFs were plated in the appropriate media (see above) in 96-well half-area black plates (Corning CLS3603) at 2500 MEFs per well for 24 hours prior to treatment. Media was exchanged for media containing doxorubicin (MilliporeSigma 324380) or etoposide (MilliporeSigma E1383) at the specified doses. Cells were incubated for the specified time. Cell viability was measured using CellTiterGlo 2.0 (Promega G9242) following manufacturer instructions.

### Statistical Analysis

Sample sizes, the number of technical or biological replicates, statistical tests, significance values, and significance thresholds are reported in the main text or figure legends pertaining to each experiment. No explicit power analysis was used to predetermine sample size. Randomization was not applicable for these experiments. No samples were excluded from analysis.

## Supplementary Text

### FLAG-IP ribosome profiling artifact

Here we present evidence that a *Tmod3* transcript detected during ribosome profiling of eS27- and eS27L-ribosomes (Figure 3D-F) is an artifact of the FLAG-IP ribosome profiling process, and not likely a *bona fide* eS27L-ribosome-associated transcript. Comparing IP RPFs to total RPFs for both the eS27-FLAG and eS27L-FLAG mESC lines, we detected that *Tmod3* mRNA was more enriched than other transcripts in the IP RPFs for both lines. This enrichment was more pronounced for the eS27L- FLAG line than for eS27-FLAG (Supplementary Figure 3A). Interestingly, Tmod3 protein is frequently reported as a contaminant of FLAG IPs in the mass spectrometry CRAPome, a repository of proteins that are non-specifically detected across many affinity purification experiments with different baits (Mellacheruvu et al., 2013). We inspected the amino acid sequence of *Tmod3* and found that it contains a sequence with high homology to the FLAG and 3xFLAG epitope tag sequences (Supplementary Figure 3B). Furthermore, we found that eS27- and eS27L-FLAG IP RPFs are relatively enriched in *Tmod3* reads at the 3’ end of the *Tmod3* coding sequence relative to the 5’ end. The beginning of this enriched interval corresponds with the base position in the coding sequence at which the ribosome would have completely translated *Tmod3’s* N-terminal FLAG-like sequence and presented it as a nascent peptide chain emerging from the ribosome’s peptide exit tunnel (Supplementary Figure 3C). We thus propose that ribosomes that are translating *Tmod3* transcripts are preferentially enriched during FLAG IP, whether or not they contain eS27-FLAG or eS27L-FLAG (Supplementary Figure 3D). We also note that the RNA concentration of IP RPFs obtained from eS27-FLAG mESCs was greater than that of eS27L-FLAG mESCs (Supplementary Figure 3E). This suggests that a larger proportion of anti-FLAG binding sites remained unoccupied during the eS27L-FLAG IP compared to the eS27-FLAG IP, and may explain why *Tmod3* appears to be more enriched among eS27L-FLAG IP RPFs.

## Supplementary Table Legends

Supplementary Table 1

Common species names, binomial nomenclature, and Uniprot IDs of the protein sequences used for multispecies alignment in Figure 1A.

Supplementary Table 2

Correlation between *eS27L* mRNA abundance (read count per 100 RP reads) and abundance of other transcripts (read count per million) in single-cell RNA-seq data (Bach et al., 2017) from alveolar cells in lactating mammary glands (‘Avd-L’ cells as termed by Bach et al.).

Supplementary Table 3

Results of paralog-specific eS27- and eS27L-FLAG ribosome profiling.

Supplementary Table 4

Organ weights and necropsy findings for 10-16-week-old mice homozygous for homogenized *eS27^eS27L^* and *eS27L^eS27^* alleles. WNL: within normal limits

Supplementary Table 5

Biomarker values for complete blood count, liver panel, and serum chemistry panel performed on 10-16-week-old homozygous males and age-matched wild-type controls from the *eS27^eS27L^* and *eS27L^eS27^* mouse lines.

Supplementary Table 6

Biomarker values for complete blood count and liver panel performed on homozygous and heterozygous 9-10-month-old males and age-matched wild-type controls from the *eS27^eS27L^* and *eS27L^eS27^* mouse lines.

Supplementary Table 7

Primer, oligonucleotide, and construct sequences used in this study.

## Source Data

Figure 3–Source Data 1: eS27 density fractionation Western blot

Figure 3–Source Data 2: eS27L density fractionation Western blot

Figure 3–Source Data 3: uS3 density fractionation Western blot

Figure 6--Source Data 1: T2018; eS27 ^+ / +^ nulliparous 8-week-old

Figure 6--Source Data 2: T2047; eS27^eS27L / +^ nulliparous 8-week-old

Figure 6--Source Data 3: T2107; eS27^eS27L / eS27L^ nulliparous 8-week-old

Figure 6--Source Data 4: T2092; eS27L^+ / +^ nulliparous 8-week-old

Figure 6--Source Data 5: T2081; eS27L^eS27 / +^ nulliparous 8-week-old

Figure 6--Source Data 6: T2051; eS27L^eS27 / eS27^ nulliparous 8-week-old

Figure 6--Source Data 7: 5x_t1850_1; eS27 ^+ / +^ pregnancy day 15

Figure 6--Source Data 8: 5x_t1866_1; eS27^eS27L / +^ pregnancy day 15

Figure 6--Source Data 9: 5x_t1931_1; eS27^eS27L / eS27L^ pregnancy day 15

Figure 6--Source Data 10: 5x_t1909_; eS27L^+ / +^ pregnancy day 15

Figure 6--Source Data 11: 5x_t1859_1; eS27L^eS27 / +^ pregnancy day 15

Figure 6--Source Data 12: 5x_t1892_1; eS27L^eS27 / eS27^ pregnancy day 15

Figure 6--Source Data 13: 5x_t915_02; eS27 ^+ / +^ lactation day 4

Figure 6--Source Data 14: 5x_t952_2; eS27^eS27L / +^ lactation day 4

Figure 6--Source Data 15: 5x_t646_3; eS27^eS27L / eS27L^ lactation day 4

Figure 6--Source Data 16: 5x_t561_02; eS27L^+ / +^ lactation day 4

Figure 6--Source Data 17: 5x_t909_; eS27L^eS27 / +^ lactation day 4

Figure 6--Source Data 18: 5x_t771_03; eS27L^eS27 / eS27^ lactation day 4

Supplementary Figure 2–Source Data 1: eS27-GFP-FLAG and eS27L-GFP-FLAG, FLAG Western blot

Supplementary Figure 2–Source Data 2: eS27-GFP-FLAG and eS27L-GFP-FLAG, eS27 Western blot

Supplementary Figure 2–Source Data 3: eS27-GFP-FLAG and eS27L-GFP-FLAG, eS27L Western blot

Supplementary Figure 2–Source Data 4: eS27-FLAG and eS27L-FLAG mESCs, eS27 Western blot

Supplementary Figure 2–Source Data 5: eS27-FLAG and eS27L-FLAG mESCs, eS27L Western blot

Supplementary Figure 2–Source Data 6: eS27-FLAG and eS27L-FLAG mESCs, FLAG Western blot

Supplementary Figure 2–Source Data 7: eS27-FLAG and eS27L-FLAG mESCs, b-actin Western blot

Supplementary Figure 2–Source Data 8: WT mESC density fractionation, FLAG Western blot

Supplementary Figure 2–Source Data 9: eS27-FLAG mESC density fractionation, FLAG Western blot

Supplementary Figure 2–Source Data 10: eS27L-FLAG mESC density fractionation, FLAG Western blot

Supplementary Figure 2–Source Data 11: eS27-FLAG and eS27L-FLAG mESC FLAG-IP, eS27 Western blot

Supplementary Figure 2–Source Data 12: eS27-FLAG and eS27L-FLAG mESC FLAG-IP, eS27L Western blot

Supplementary Figure 2–Source Data 13: eS27-FLAG and eS27L-FLAG mESC FLAG-IP, uS3 Western blot

Supplementary Figure 2–Source Data 14: eS27-FLAG and eS27L-FLAG mESC FLAG-IP, GAPDH Western blot

## Acknowledgements

We thank members of the Barna, Pritchard, and Attardi labs for advice and critiques, Yiran Liu for advice on designing CRISPR strategies, Elias Godoy for assistance with necropsy dissections, the Stanford Animal Histology Core for assistance with histology sample preparation, and the Stanford Animal Diagnostic lab for preparation of histology samples. We thank the Stanford Shared FACS Facility for assistance with flow cytometry experiments. We thank the UCSF Gladstone Institutes Transgenic Gene-Targeting Core for facility support in generating CRISPR-edited mice. pSpCas9(BB)-2A-Puro (PX459) V2.0 was a gift from Feng Zhang (Addgene plasmid # 62988 ; http://n2t.net/addgene:62988 ; RRID:Addgene_62988)

## Author contributions

A.X., J.K.P., and M.B. conceived the project. A.X., R.M., E.F., H.T., Z.Z., J.M., and K.C. planned, performed, and analyzed the experiments. D.R. assisted with the generation of CRISPR- edited mouse lines. L.H. advised on mammary gland experiments. A.X. wrote the manuscript in consultation with J.K.P. and M.B. and with assistance from R.M. and K.C. All authors have reviewed, critiqued, and approved the manuscript.

## Supplementary Figure Legends

**Supplementary Figure 1 related to Figure 1.**
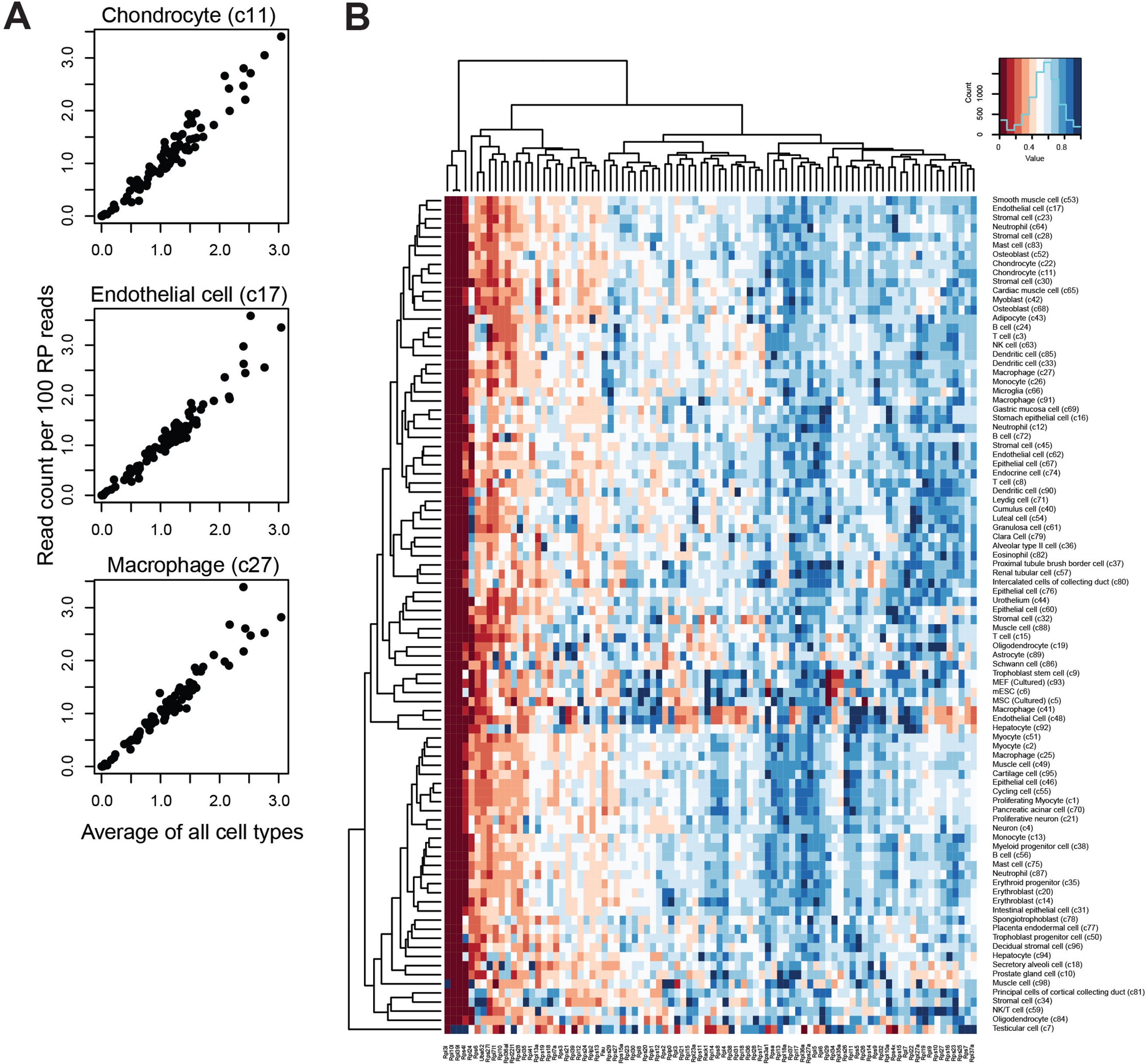
**(A)** Using single-cell RNA-seq data from the Mouse Cell Atlas (Han et al., 2018), correlation of transcript abundance among RP genes is plotted for three example cell types (y-axis) against the average values across all cell types (x-axis). Each point represents an RP gene. Chondrocytes: Pearson’s r = 0.97, p-value < 2.2e-16. Endothelial cells: Pearson’s r = 0.97, p-value < 2.2e-16. Macrophages: Pearson’s r = 0.97, p-value < 2.2e-16. “c11”, “c17”, and “c27” refer to the cell type cluster IDs as annotated by Han et al. **(B)** Expression patterns for RP genes across all cell types in the Mouse Cell Atlas. Read counts are normalized per 100 RP reads and scaled to the maximum value for each gene.

**Supplementary Figure 2 related to Figure 3.**
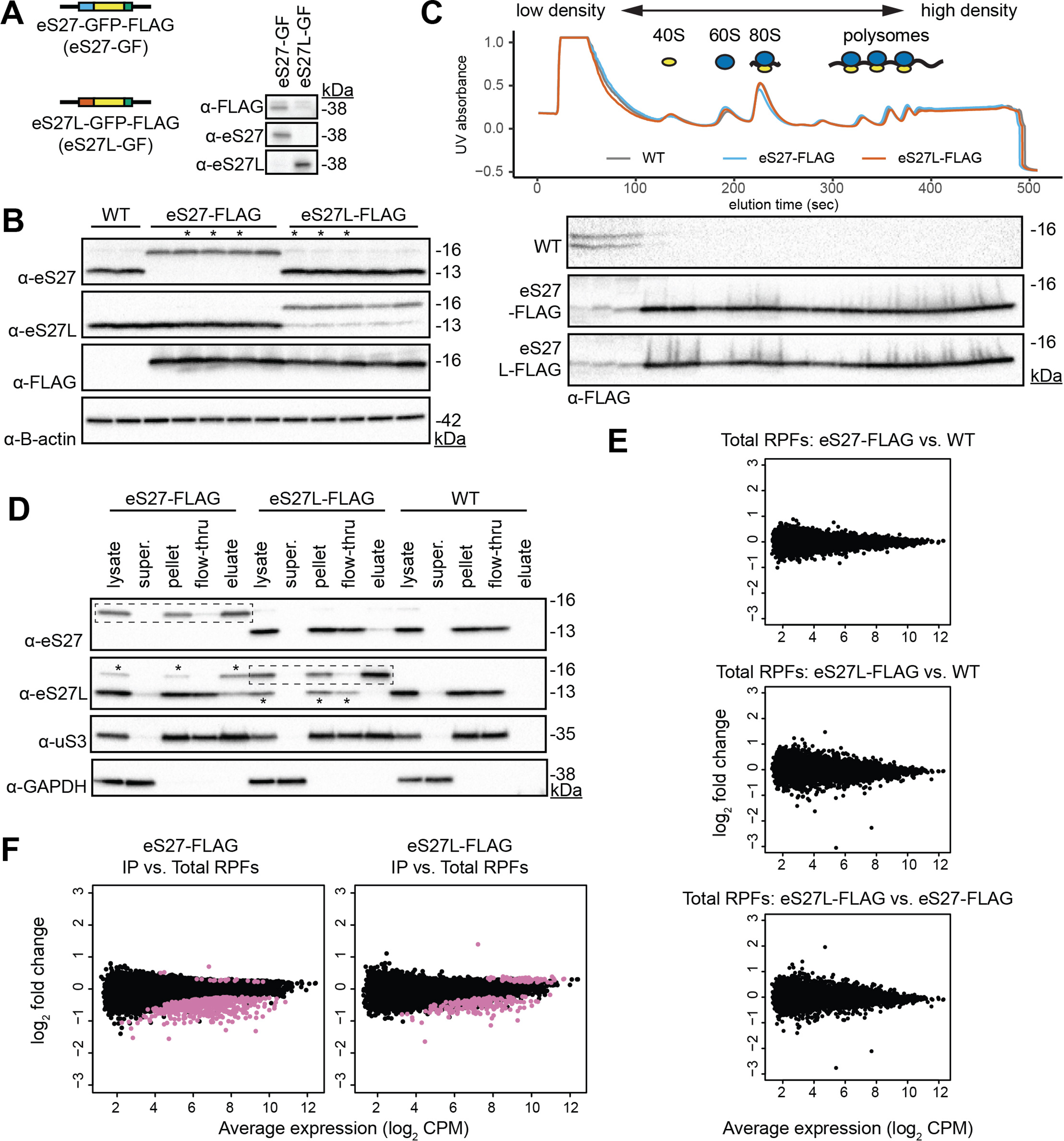
**(A)** Plasmids encoding eS27- and eS27L-GFP fusion proteins were transfected into mouse embryonic stem cells (mESCs). A Western blot was performed on cell lysate using commercially available antibodies to assess whether they are paralog-specific. GFP fusion adds 27 kilo-Daltons (kDa) of molecular weight. The anti-eS27L antibody has trace detection of eS27. **(B)** Western blots of cell lysates from WT, eS27-FLAG and eS27L-FLAG mESCs. Each lane is an independently selected clone. Beta actin is used as a loading control. Clones with an asterisk (*) were used as biological replicates in subsequent experiments. **(C)** Western blot of density-fractionated cell lysates from eS27- and eS27L-FLAG mESCs. A representative UV absorbance trace and blot from each mESC line is shown. **(D)** Western blot of cell lysates from one biological replicate each of eS27- FLAG, eS27L-FLAG, and WT mESCs; supernatant containing no ribosomes after ultracentrifugation; pellet containing total ribosomes after centrifugation, used as input for FLAG immunoprecipitation (IP); flow-through of FLAG IP, and eluate of FLAG-IP. uS3 is shown as a typical ribosomal protein (RP) that is also in the 40S small subunit. GAPDH is shown as a typical protein that is not ribosome-associat- ed. Dashed boxes indicate the eS27-FLAG or eS27L-FLAG protein targeted for IP in the respective cell lines. Asterisks in the anti-eS- 27L blot indicate trace detection of eS27 by the eS27L antibody. **(E)** Comparison of total ribosome-protected fragments (RPFs) between eS27-FLAG, eS27L-FLAG and passage-matched wild-type (WT) control cell line. CPM = counts per million. **(F)** Comparison of IP RPFs versus total RPFs in eS27-FLAG and eS27L-FLAG mESCs. Purple indicates enriched or depleted transcripts at a multiple-hypothesis-corrected false discovery rate (FDR) <0.05, using the same statistical test described in Figure 3. See also Supplementary Table 3 and Supplementary Figure 2--Source Data 1-14.

**Supplementary Figure 3 related to Figure 3.**
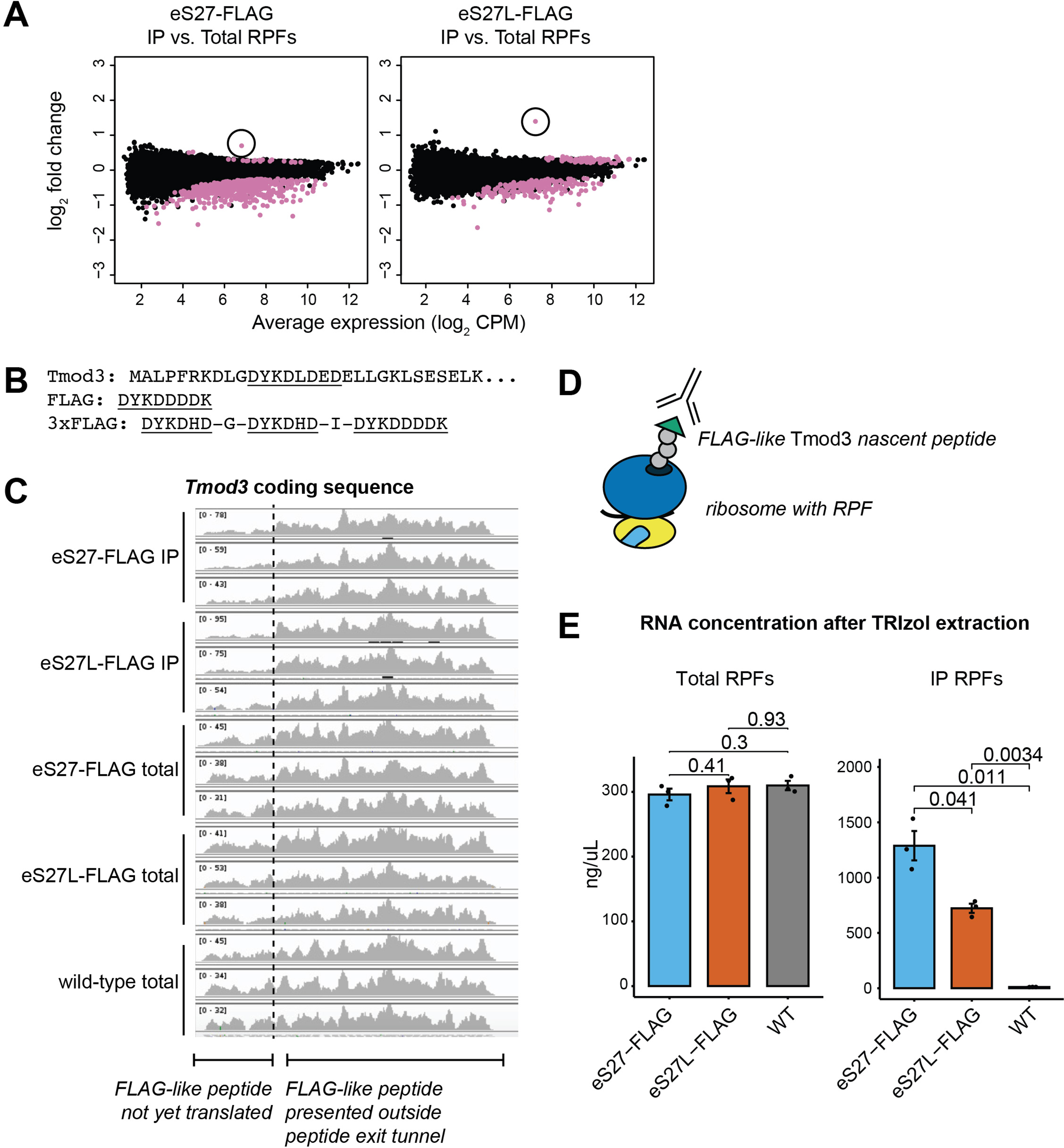
**(A)** Enrichment of *Tmod3* mRNA (circled) among immunoprecipitated ribosome-protected fragments (IP RPFs) from both eS27- and eS27L-FLAG mESCs, relative to total RPFs. **(B)** N-terminal sequence of *Tmod3* protein, with FLAG-like peptide sequence highlighted. **(C)** Distribution of IP RPFs and total RPFs along the coding sequence of *Tmod3*. Base positions encoding the FLAG-like peptide are denoted with an arrow. Note that the y-axis range differs for each sample. **(D)** Proposed binding of the anti-FLAG antibody to the FLAG-like *Tmod3* nascent peptide. **(E)** RNA concentration as estimated by absor- bance at 260 and 280 nm wavelength after TRIzol extraction of total RPF and IP RPF samples from eS27-FLAG, eS27L-FLAG, and WT mESCs, showing consistently lower IP yield of eS27L-ribosomes compared to eS27L-ribosomes. Significance was assessed by t-test. n=3 biological replicates.

**Supplementary Figure 4 related to Figure 5.**
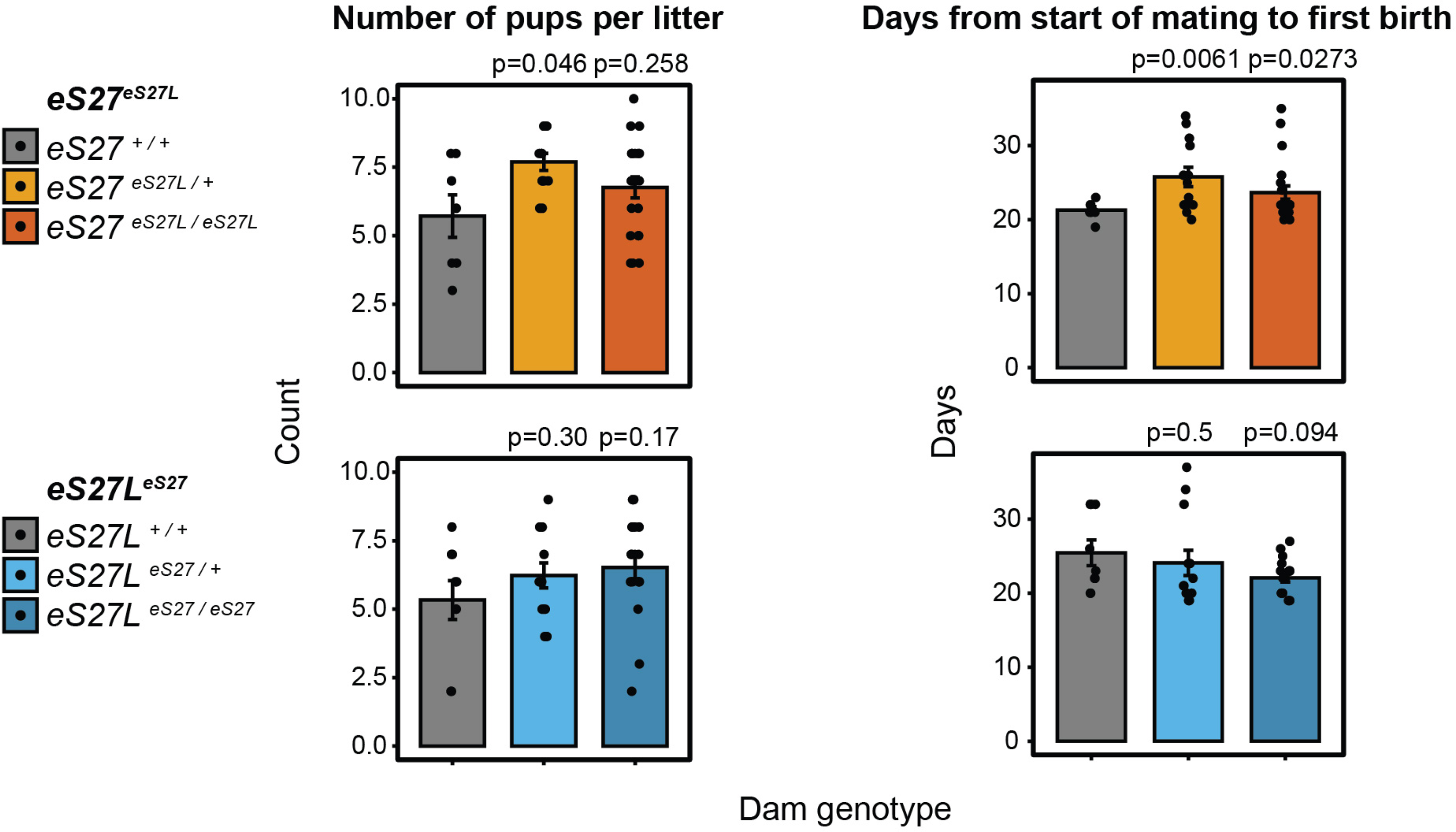
Comparison of litter size (number of pups) and days between start of mating and first birth for all dam genotypes of homogenized *eS27eS27L* and *eS27LeS27* lineages. n = 5-19 litters per dam genotype and timepoint. Significance relative to WT was assessed by t-test. All females were housed with stud males beginning at 8 weeks of age. Only first litters are included. Statistically significant differences in litter size and mating-birth interval were observed in *eS27 ^eS27L / +^* dams relative to *eS27 ^+ / +^* dams, and a statistically significant difference in mating-birth interval was observed in *eS27 ^eS27L / eS27L^* dams relative to *eS27 ^+ / +^* dams. However, considering the litter sizes and mating-birth intervals observed in *eS27L ^+ / +^* dams, we conclude that these variations are within the normal phenotypic range for wild-type animals.

## References

Anger, A. M., Armache, J.-P., Berninghausen, O., Habeck, M., Subklewe, M., Wilson, D. N., & Beckmann, R. (2013). Structures of the human and Drosophila 80S ribosome. Nature, 497(7447), 80–85. https://doi.org/10.1038/nature12104

Antebi, Y. E., Linton, J. M., Klumpe, H., Bintu, B., Gong, M., Su, C., McCardell, R., & Elowitz, M. B. (2017). Combinatorial signal perception in the BMP pathway. Cell, 170(6), 1184–1196.e24. https://doi.org/10.1016/j.cell.2017.08.015

Ashcroft, M., & Vousden, K. H. (1999). Regulation of p53 stability. Oncogene, 18(53), 7637–7643. https://doi.org/10.1038/sj.onc.1203012

Aury, J.-M., Jaillon, O., Duret, L., Noel, B., Jubin, C., Porcel, B. M., Ségurens, B., Daubin, V., Anthouard, V., Aiach, N., Arnaiz, O., Billaut, A., Beisson, J., Blanc, I., Bouhouche, K., Câmara, F., Duharcourt, S., Guigo, R., Gogendeau, D., … Wincker, P. (2006). Global trends of whole-genome duplications revealed by the ciliate *Paramecium tetraurelia*. Nature, 444(7116), 171–178. https://doi.org/10.1038/nature05230

Bach, K., Pensa, S., Grzelak, M., Hadfield, J., Adams, D. J., Marioni, J. C., & Khaled, W. T. (2017). Differentiation dynamics of mammary epithelial cells revealed by single-cell RNA sequencing. Nature Communications, 8(1), 2128. https://doi.org/10.1038/s41467-017-02001-5

Balasubramanian, S., Zheng, D., Liu, Y.-J., Fang, G., Frankish, A., Carriero, N., Robilotto, R., Cayting, P., & Gerstein, M. (2009). Comparative analysis of processed ribosomal protein pseudogenes in four mammalian genomes. Genome Biology, 10(1), R2. https://doi.org/10.1186/gb-2009-10-1-r2

Bruce, A. E., Oates, A. C., Prince, V. E., & Ho, R. K. (2001). Additional hox clusters in the zebrafish: divergent expression patterns belie equivalent activities of duplicate hoxB5 genes. Evolution & Development, 3(3), 127–144. https://doi.org/10.1046/j.1525-142x.2001.003003127.x

Chaillou, T., Zhang, X., & McCarthy, J. J. (2016). Expression of Muscle-Specific Ribosomal Protein L3- Like Impairs Myotube Growth. Journal of Cellular Physiology, 231(9), 1894–1902. https://doi.org/10.1002/jcp.25294

Conant, G. C., & Wolfe, K. H. (2006). Functional partitioning of yeast co-expression networks after genome duplication. PLoS Biology, 4(4), e109. https://doi.org/10.1371/journal.pbio.0040109

Conant, G. C., & Wolfe, K. H. (2008). Turning a hobby into a job: how duplicated genes find new functions. Nature Reviews. Genetics, 9(12), 938–950. https://doi.org/10.1038/nrg2482

Dehal, P., & Boore, J. L. (2005). Two rounds of whole genome duplication in the ancestral vertebrate. PLoS Biology, 3(10), e314. https://doi.org/10.1371/journal.pbio.0030314

De Kegel, B., & Ryan, C. J. (2019). Paralog buffering contributes to the variable essentiality of genes in cancer cell lines. PLoS Genetics, 15(10), e1008466. https://doi.org/10.1371/journal.pgen.1008466

Doench, J. G., Fusi, N., Sullender, M., Hegde, M., Vaimberg, E. W., Donovan, K. F., Smith, I., Tothova, Z., Wilen, C., Orchard, R., Virgin, H. W., Listgarten, J., & Root, D. E. (2016). Optimized sgRNA design to maximize activity and minimize off-target effects of CRISPR-Cas9. Nature Biotechnology, 34(2), 184–191. https://doi.org/10.1038/nbt.3437

Durkin, M. E., Qian, X., Popescu, N. C., & Lowy, D. R. (2013). Isolation of Mouse Embryo Fibroblasts. Bio-Protocol, 3(18). https://doi.org/10.21769/BioProtoc.908

Force, A., Lynch, M., Pickett, F. B., Amores, A., Yan, Y. L., & Postlethwait, J. (1999). Preservation of duplicate genes by complementary, degenerative mutations. Genetics, 151(4), 1531–1545. https://doi.org/10.1093/genetics/151.4.1531

Fu, N. Y., Rios, A. C., Pal, B., Soetanto, R., Lun, A. T. L., Liu, K., Beck, T., Best, S. A., Vaillant, F., Bouillet, P., Strasser, A., Preiss, T., Smyth, G. K., Lindeman, G. J., & Visvader, J. E. (2015). EGF- mediated induction of Mcl-1 at the switch to lactation is essential for alveolar cell survival. Nature Cell Biology, 17(4), 365–375. https://doi.org/10.1038/ncb3117

Genuth, N. R., & Barna, M. (2018). Heterogeneity and specialized functions of translation machinery: from genes to organisms. Nature Reviews. Genetics, 19(7), 431–452. https://doi.org/10.1038/s41576-018-0008-z

Gerst, J. E. (2018). Pimp my ribosome: ribosomal protein paralogs specify translational control. Trends in Genetics, 34(11), 832–845. https://doi.org/10.1016/j.tig.2018.08.004

Ghulam, M. M., Catala, M., & Abou Elela, S. (2020). Differential expression of duplicated ribosomal protein genes modifies ribosome composition in response to stress. Nucleic Acids Research, 48(4), 1954–1968. https://doi.org/10.1093/nar/gkz1183

Graur, D., & Li, W.-H. (1997). Fundamentals of Molecular Evolution (2nd ed.). https://www.sinauer.com/media/wysiwyg/tocs/FundamentalsMolecularEvolution.pdf

Greer, J. M., Puetz, J., Thomas, K. R., & Capecchi, M. R. (2000). Maintenance of functional equivalence during paralogous Hox gene evolution. Nature, 403(6770), 661–665. https://doi.org/10.1038/35001077

Guimaraes, J. C., & Zavolan, M. (2016). Patterns of ribosomal protein expression specify normal and malignant human cells. Genome Biology, 17(1), 236. https://doi.org/10.1186/s13059-016-1104-z

Gupta, V., & Warner, J. R. (2014). Ribosome-omics of the human ribosome. RNA (New York), 20(7), 1004–1013. https://doi.org/10.1261/rna.043653.113

Gu, Z., Steinmetz, L. M., Gu, X., Scharfe, C., Davis, R. W., & Li, W.-H. (2003). Role of duplicate genes in genetic robustness against null mutations. Nature, 421(6918), 63–66. https://doi.org/10.1038/nature01198

Han, X., Wang, R., Zhou, Y., Fei, L., Sun, H., Lai, S., Saadatpour, A., Zhou, Z., Chen, H., Ye, F., Huang, D., Xu, Y., Huang, W., Jiang, M., Jiang, X., Mao, J., Chen, Y., Lu, C., Xie, J., … Guo, G. (2018). Mapping the Mouse Cell Atlas by Microwell-Seq. Cell, 172(5), 1091–1107.e17. https://doi.org/10.1016/j.cell.2018.02.001

He, H., & Sun, Y. (2007). Ribosomal protein S27L is a direct p53 target that regulates apoptosis. Oncogene, 26(19), 2707–2716. https://doi.org/10.1038/sj.onc.1210073

Hsu, P. D., Scott, D. A., Weinstein, J. A., Ran, F. A., Konermann, S., Agarwala, V., Li, Y., Fine, E. J., Wu, X., Shalem, O., Cradick, T. J., Marraffini, L. A., Bao, G., & Zhang, F. (2013). DNA targeting specificity of RNA-guided Cas9 nucleases. Nature Biotechnology, 31(9), 827–832. https://doi.org/10.1038/nbt.2647

Hughes, A. L. (1994). The evolution of functionally novel proteins after gene duplication. Proceedings. Biological Sciences / the Royal Society, 256(1346), 119–124. https://doi.org/10.1098/rspb.1994.0058

Ingolia, N. T., Ghaemmaghami, S., Newman, J. R. S., & Weissman, J. S. (2009). Genome-wide analysis in vivo of translation with nucleotide resolution using ribosome profiling. Science, 324(5924), 218–223. https://doi.org/10.1126/science.1168978

Innan, H., & Kondrashov, F. (2010). The evolution of gene duplications: classifying and distinguishing between models. Nature Reviews. Genetics, 11(2), 97–108. https://doi.org/10.1038/nrg2689

Jiang, L., Li, T., Zhang, X., Zhang, B., Yu, C., Li, Y., Fan, S., Jiang, X., Khan, T., Hao, Q., Xu, P., Nadano, D., Huleihel, M., Lunenfeld, E., Wang, P. J., Zhang, Y., & Shi, Q. (2017). RPL10L Is Required for Male Meiotic Division by Compensating for RPL10 during Meiotic Sex Chromosome Inactivation in Mice. Current Biology, 27(10), 1498–1505.e6. https://doi.org/10.1016/j.cub.2017.04.017

Kao, B. R., Malerba, A., Lu-Nguyen, N. B., Harish, P., McCarthy, J. J., Dickson, G., & Popplewell, L. J. (2021). Knockdown of Muscle-Specific Ribosomal Protein L3-Like Enhances Muscle Function in Healthy and Dystrophic Mice. Nucleic Acid Therapeutics, 31(6), 457–464. https://doi.org/10.1089/nat.2020.0928

Komili, S., Farny, N. G., Roth, F. P., & Silver, P. A. (2007). Functional specificity among ribosomal proteins regulates gene expression. Cell, 131(3), 557–571. https://doi.org/10.1016/j.cell.2007.08.037

Kondrashov, F. A., Rogozin, I. B., Wolf, Y. I., & Koonin, E. V. (2002). Selection in the evolution of gene duplications. Genome Biology, 3(2), RESEARCH0008. https://doi.org/10.1186/gb-2002-3-2-research0008

Kuang, G., Tao, W., Zheng, S., Wang, X., & Wang, D. (2020). Genome-Wide Identification, Evolution and Expression of the Complete Set of Cytoplasmic Ribosomal Protein Genes in Nile Tilapia. International Journal of Molecular Sciences, 21(4). https://doi.org/10.3390/ijms21041230

Langmead, B., & Salzberg, S. L. (2012). Fast gapped-read alignment with Bowtie 2. Nature Methods, 9(4), 357–359. https://doi.org/10.1038/nmeth.1923

Lan, X., & Pritchard, J. K. (2016). Coregulation of tandem duplicate genes slows evolution of subfunctionalization in mammals. Science, 352(6288), 1009–1013. https://doi.org/10.1126/science.aad8411

Law, C. W., Chen, Y., Shi, W., & Smyth, G. K. (2014). voom: Precision weights unlock linear model analysis tools for RNA-seq read counts. Genome Biology, 15(2), R29. https://doi.org/10.1186/gb-2014-15-2-r29

Li, J., Tan, J., Zhuang, L., Banerjee, B., Yang, X., Chau, J. F. L., Lee, P. L., Hande, M. P., Li, B., & Yu, Q. (2007). Ribosomal protein S27-like, a p53-inducible modulator of cell fate in response to genotoxic stress. Cancer Research, 67(23), 11317–11326. https://doi.org/10.1158/0008-5472.CAN-07-1088

Lynch, M., & Conery, J. S. (2000). The evolutionary fate and consequences of duplicate genes. Science, 290(5494), 1151–1155. https://doi.org/10.1126/science.290.5494.1151

Lynch, M. (2007). The frailty of adaptive hypotheses for the origins of organismal complexity. Proceedings of the National Academy of Sciences of the United States of America, 104 *Suppl 1*, 8597–8604. https://doi.org/10.1073/pnas.0702207104

Macias, H., & Hinck, L. (2012). Mammary gland development. Wiley Interdisciplinary Reviews. Developmental Biology, 1(4), 533–557. https://doi.org/10.1002/wdev.35

Makino, T., & McLysaght, A. (2010). Ohnologs in the human genome are dosage balanced and frequently associated with disease. Proceedings of the National Academy of Sciences of the United States of America, 107(20), 9270–9274. https://doi.org/10.1073/pnas.0914697107

Manchado, M., Infante, C., Asensio, E., Cañavate, J. P., & Douglas, S. E. (2007). Comparative sequence analysis of the complete set of 40S ribosomal proteins in the Senegalese sole (Solea senegalensis Kaup) and Atlantic halibut (Hippoglossus hippoglossus L.) (Teleostei: Pleuronectiformes): phylogeny and tissue- and development-specific expression. BMC Evolutionary Biology, 7, 107. https://doi.org/10.1186/1471-2148-7-107

Martin, M. (2011). Cutadapt removes adapter sequences from high-throughput sequencing reads. EMBnet.Journal, 17(1), 10. https://doi.org/10.14806/ej.17.1.200

McGlincy, N. J., & Ingolia, N. T. (2017). Transcriptome-wide measurement of translation by ribosome profiling. Methods, 126, 112–129. https://doi.org/10.1016/j.ymeth.2017.05.028

Meek, D. W., & Anderson, C. W. (2009). Posttranslational modification of p53: cooperative integrators of function. Cold Spring Harbor Perspectives in Biology, 1(6), a000950. https://doi.org/10.1101/cshperspect.a000950

Mellacheruvu, D., Wright, Z., Couzens, A. L., Lambert, J.-P., St-Denis, N. A., Li, T., Miteva, Y. V., Hauri, S., Sardiu, M. E., Low, T. Y., Halim, V. A., Bagshaw, R. D., Hubner, N. C., Al-Hakim, A., Bouchard, A., Faubert, D., Fermin, D., Dunham, W. H., Goudreault, M., … Nesvizhskii, A. I. (2013). The CRAPome: a contaminant repository for affinity purification-mass spectrometry data. Nature Methods, 10(8), 730–736. https://doi.org/10.1038/nmeth.2557

Milenkovic, I., Santos Vieira, H. G., Lucas, M. C., Ruiz-Orera, J., Patone, G., Kesteven, S., Wu, J., Feneley, M., Espadas, G., Sabidó, E., Hubner, N., van Heesch, S., Voelkers, M., & Novoa, E. M. (2021). RPL3L-containing ribosomes modulate mitochondrial activity in the mammalian heart. BioRxiv. https://doi.org/10.1101/2021.12.04.471171

Nadeau, J. H., & Sankoff, D. (1997). Comparable rates of gene loss and functional divergence after genome duplications early in vertebrate evolution. Genetics, 147(3), 1259–1266.

Nakao, A., Yoshihama, M., & Kenmochi, N. (2004). RPG: the Ribosomal Protein Gene database. Nucleic Acids Research, 32(Database issue), D168-70. https://doi.org/10.1093/nar/gkh004

Natchiar, S. K., Myasnikov, A. G., Kratzat, H., Hazemann, I., & Klaholz, B. P. (2017). Visualization of chemical modifications in the human 80S ribosome structure. Nature, 551(7681), 472–477. https://doi.org/10.1038/nature24482

Nicolas, E., Parisot, P., Pinto-Monteiro, C., de Walque, R., De Vleeschouwer, C., & Lafontaine, D. L. J. (2016). Involvement of human ribosomal proteins in nucleolar structure and p53-dependent nucleolar stress. Nature Communications, 7, 11390. https://doi.org/10.1038/ncomms11390

O’Donohue, M.-F., Choesmel, V., Faubladier, M., Fichant, G., & Gleizes, P.-E. (2010). Functional dichotomy of ribosomal proteins during the synthesis of mammalian 40S ribosomal subunits. The Journal of Cell Biology, 190(5), 853–866. https://doi.org/10.1083/jcb.201005117

Ohno, S. (1970). Evolution by gene duplication. Springer Berlin Heidelberg. https://doi.org/10.1007/978-3-642-86659-3

Palmer, C. A., Neville, M. C., Anderson, S. M., & McManaman, J. L. (2006). Analysis of lactation defects in transgenic mice. Journal of Mammary Gland Biology and Neoplasia, 11(3–4), 269–282. https://doi.org/10.1007/s10911-006-9023-3

Papp, B., Pál, C., & Hurst, L. D. (2003). Dosage sensitivity and the evolution of gene families in yeast. Nature, 424(6945), 194–197. https://doi.org/10.1038/nature01771

Parenteau, J., Lavoie, M., Catala, M., Malik-Ghulam, M., Gagnon, J., & Abou Elela, S. (2015). Preservation of Gene Duplication Increases the Regulatory Spectrum of Ribosomal Protein Genes and Enhances Growth under Stress. Cell Reports, 13(11), 2516–2526. https://doi.org/10.1016/j.celrep.2015.11.033

Patrinostro, X., Roy, P., Lindsay, A., Chamberlain, C. M., Sundby, L. J., Starker, C. G., Voytas, D. F., Ervasti, J. M., & Perrin, B. J. (2018). Essential nucleotide- and protein-dependent functions of Actb/β-actin. Proceedings of the National Academy of Sciences of the United States of America, 115(31), 7973–7978. https://doi.org/10.1073/pnas.1807895115

Plante, I., Stewart, M. K. G., & Laird, D. W. (2011). Evaluation of mammary gland development and function in mouse models. Journal of Visualized Experiments, 53. https://doi.org/10.3791/2828

Prince, V. E., & Pickett, F. B. (2002). Splitting pairs: the diverging fates of duplicated genes. Nature Reviews. Genetics, 3(11), 827–837. https://doi.org/10.1038/nrg928

Raices, M., & D’Angelo, M. A. (2012). Nuclear pore complex composition: a new regulator of tissue- specific and developmental functions. Nature Reviews. Molecular Cell Biology, 13(11), 687–699. https://doi.org/10.1038/nrm3461

Ran, F. A., Hsu, P. D., Wright, J., Agarwala, V., Scott, D. A., & Zhang, F. (2013). Genome engineering using the CRISPR-Cas9 system. Nature Protocols, 8(11), 2281–2308. https://doi.org/10.1038/nprot.2013.143

Ritchie, M. E., Phipson, B., Wu, D., Hu, Y., Law, C. W., Shi, W., & Smyth, G. K. (2015). limma powers differential expression analyses for RNA-sequencing and microarray studies. Nucleic Acids Research, 43(7), e47. https://doi.org/10.1093/nar/gkv007

Robinson, M. D., McCarthy, D. J., & Smyth, G. K. (2010). edgeR: a Bioconductor package for differential expression analysis of digital gene expression data. Bioinformatics, 26(1), 139–140. https://doi.org/10.1093/bioinformatics/btp616

Russo, A., & Russo, G. (2017). Ribosomal Proteins Control or Bypass p53 during Nucleolar Stress. International Journal of Molecular Sciences, 18(1). https://doi.org/10.3390/ijms18010140

Sacerdot, C., Louis, A., Bon, C., Berthelot, C., & Roest Crollius, H. (2018). Chromosome evolution at the origin of the ancestral vertebrate genome. Genome Biology, 19(1), 166. https://doi.org/10.1186/s13059-018-1559-1

Sidow, A. (1996). Gen(om)e duplications in the evolution of early vertebrates. Current Opinion in Genetics & Development, 6(6), 715–722. https://doi.org/10.1016/S0959-437X(96)80026-8

Singh, P. P., & Isambert, H. (2020). OHNOLOGS v2: a comprehensive resource for the genes retained from whole genome duplication in vertebrates. Nucleic Acids Research, 48(D1), D724–D730. https://doi.org/10.1093/nar/gkz909

Smith, T., Heger, A., & Sudbery, I. (2017). UMI-tools: modeling sequencing errors in Unique Molecular Identifiers to improve quantification accuracy. Genome Research, 27(3), 491–499. https://doi.org/10.1101/gr.209601.116

Stark, G. R., & Wahl, G. M. (1984). Gene amplification. Annual Review of Biochemistry, 53, 447–491. https://doi.org/10.1146/annurev.bi.53.070184.002311

Sugihara, Y., Honda, H., Iida, T., Morinaga, T., Hino, S., Okajima, T., Matsuda, T., & Nadano, D. (2010). Proteomic analysis of rodent ribosomes revealed heterogeneity including ribosomal proteins L10-like, L22-like 1, and L39-like. Journal of Proteome Research, 9(3), 1351–1366. https://doi.org/10.1021/pr9008964

Sun, S., He, H., Ma, Y., Xu, J., Chen, G., Sun, Y., & Xiong, X. (2020). Inactivation of ribosomal protein S27-like impairs DNA interstrand cross-link repair by destabilization of FANCD2 and FANCI. Cell Death & Disease, 11(10), 852. https://doi.org/10.1038/s41419-020-03082-9

Tabula Muris Consortium, Overall coordination, Logistical coordination, Organ collection and processing, Library preparation and sequencing, Computational data analysis, Cell type annotation, Writing group, Supplemental text writing group, & Principal investigators. (2018). Single-cell transcriptomics of 20 mouse organs creates a Tabula Muris. Nature, 562(7727), 367– 372. https://doi.org/10.1038/s41586-018-0590-4

Taggart, J. C., Zauber, H., Selbach, M., Li, G.-W., & McShane, E. (2020). Keeping the proportions of protein complex components in check. Cell Systems, 10(2), 125–132. https://doi.org/10.1016/j.cels.2020.01.004

Thompson, N. A., Ranzani, M., van der Weyden, L., Iyer, V., Offord, V., Droop, A., Behan, F., Gonçalves, E., Speak, A., Iorio, F., Hewinson, J., Harle, V., Robertson, H., Anderson, E., Fu, B., Yang, F., Zagnoli-Vieira, G., Chapman, P., Del Castillo Velasco-Herrera, M., … Adams, D. J. (2021). Combinatorial CRISPR screen identifies fitness effects of gene paralogues. Nature Communications, 12(1), 1302. https://doi.org/10.1038/s41467-021-21478-9

Topisirovic, I., & Sonenberg, N. (2011). Translational control by the eukaryotic ribosome. Cell, 145(3), 333–334. https://doi.org/10.1016/j.cell.2011.04.006

Uechi, T., Tanaka, T., & Kenmochi, N. (2001). A complete map of the human ribosomal protein genes: assignment of 80 genes to the cytogenetic map and implications for human disorders. Genomics, 72(3), 223–230. https://doi.org/10.1006/geno.2000.6470

Vedula, P., Kurosaka, S., Leu, N. A., Wolf, Y. I., Shabalina, S. A., Wang, J., Sterling, S., Dong, D. W., & Kashina, A. (2017). Diverse functions of homologous actin isoforms are defined by their nucleotide, rather than their amino acid sequence. ELife, 6. https://doi.org/10.7554/eLife.31661

Wapinski, I., Pfeffer, A., Friedman, N., & Regev, A. (2007). Natural history and evolutionary principles of gene duplication in fungi. Nature, 449(7158), 54–61. https://doi.org/10.1038/nature06107

Warner, J. R., & McIntosh, K. B. (2009). How common are extraribosomal functions of ribosomal proteins? Molecular Cell, 34(1), 3–11. https://doi.org/10.1016/j.molcel.2009.03.006

Wong, Q. W.-L., Li, J., Ng, S. R., Lim, S. G., Yang, H., & Vardy, L. A. (2014). RPL39L is an example of a recently evolved ribosomal protein paralog that shows highly specific tissue expression patterns and is upregulated in ESCs and HCC tumors. RNA Biology, 11(1), 33–41. https://doi.org/10.4161/rna.27427

Xiong, X., Cui, D., Bi, Y., Sun, Y., & Zhao, Y. (2020). Neddylation modification of ribosomal protein RPS27L or RPS27 by MDM2 or NEDP1 regulates cancer cell survival. The FASEB Journal, 34(10), 13419–13429. https://doi.org/10.1096/fj.202000530RRR

Xiong, X., Liu, X., Li, H., He, H., Sun, Y., & Zhao, Y. (2018). Ribosomal protein S27-like regulates autophagy via the β-TrCP-DEPTOR-mTORC1 axis. Cell Death & Disease, 9(11), 1131. https://doi.org/10.1038/s41419-018-1168-7

Xiong, X., Zhao, Y., He, H., & Sun, Y. (2011). Ribosomal protein S27-like and S27 interplay with p53- MDM2 axis as a target, a substrate and a regulator. Oncogene, 30(15), 1798–1811. https://doi.org/10.1038/onc.2010.569

Xiong, X., Zhao, Y., Tang, F., Wei, D., Thomas, D., Wang, X., Liu, Y., Zheng, P., & Sun, Y. (2014). Ribosomal protein S27-like is a physiological regulator of p53 that suppresses genomic instability and tumorigenesis. ELife, 3, e02236. https://doi.org/10.7554/eLife.02236

Xue, S., & Barna, M. (2012). Specialized ribosomes: a new frontier in gene regulation and organismal biology. Nature Reviews. Molecular Cell Biology, 13(6), 355–369. https://doi.org/10.1038/nrm3359

Zhang, J.-P., Li, X.-L., Li, G.-H., Chen, W., Arakaki, C., Botimer, G. D., Baylink, D., Zhang, L., Wen, W., Fu, Y.-W., Xu, J., Chun, N., Yuan, W., Cheng, T., & Zhang, X.-B. (2017). Efficient precise knockin with a double cut HDR donor after CRISPR/Cas9-mediated double-stranded DNA cleavage. Genome Biology, 18(1), 35. https://doi.org/10.1186/s13059-017-1164-8

Zhang, Yanping, & Lu, H. (2009). Signaling to p53: ribosomal proteins find their way. Cancer Cell, 16(5), 369–377. https://doi.org/10.1016/j.ccr.2009.09.024

Zhang, Yong, O’Leary, M. N., Peri, S., Wang, M., Zha, J., Melov, S., Kappes, D. J., Feng, Q., Rhodes, J., Amieux, P. S., Morris, D. R., Kennedy, B. K., & Wiest, D. L. (2017). Ribosomal Proteins Rpl22 and Rpl22l1 Control Morphogenesis by Regulating Pre-mRNA Splicing. Cell Reports, 18(2), 545– 556. https://doi.org/10.1016/j.celrep.2016.12.034

Zhao, Y., Tan, M., Liu, X., Xiong, X., & Sun, Y. (2018). Inactivation of ribosomal protein S27-like confers radiosensitivity via the Mdm2-p53 and Mdm2-MRN-ATM axes. Cell Death & Disease, 9(2), 145. https://doi.org/10.1038/s41419-017-0192-3

Zou, Q., & Qi, H. (2021). Deletion of ribosomal paralogs Rpl39 and Rpl39l compromises cell proliferation via protein synthesis and mitochondrial activity. The International Journal of Biochemistry & Cell Biology, 139, 106070. https://doi.org/10.1016/j.biocel.2021.106070

Zou, Q., Yang, L., Shi, R., Qi, Y., Zhang, X., & Qi, H. (2021). Proteostasis regulated by testis-specific ribosomal protein RPL39L maintains mouse spermatogenesis. IScience, 24(12), 103396. https://doi.org/10.1016/j.isci.2021.103396

